# Expression of a constitutively active nitrate reductase increases SARS-CoV-2 Spike protein production in *Nicotiana benthamiana* leaves that otherwise show traits of senescence

**DOI:** 10.1101/2024.09.23.614552

**Authors:** Louis-Philippe Hamel, Marie-Ève Paré, Francis Poirier-Gravel, Rachel Tardif, Marc-André Comeau, Pierre-Olivier Lavoie, Andréane Langlois, Marie-Claire Goulet, Marc-André D’Aoust, Dominique Michaud

## Abstract

The production of coronavirus disease 2019 vaccines can be achieved by transient expression of the Spike (S) protein of Severe Acute Respiratory Syndrome Coronavirus 2 in agroinfiltrated leaves of *Nicotiana benthamiana*, a process promoted by the co-expression of viral silencing suppressor P19. Upon expression, the S protein enters the cell secretory pathway, before being trafficked to the plasma membrane where formation of coronavirus-like particles (CoVLPs) occurs. We recently used RNAseq and time course sampling to characterize molecular responses of *N. benthamiana* leaf cells expressing P19 only, or P19 in combination with recombinant S protein. This revealed expression of the viral proteins to deeply affect the physiological status of plant cells, including through the activation of immune responses. Here, transcriptomics shows that the production of CoVLPs also induces leaf senescence, as revealed by the upregulation of senescence-associated genes, activation of senescence-related proteases, and downregulation of genes involved in basic metabolic functions like photosynthesis or nitrogen uptake and assimilation. CoVLP production also upregulated asparagine synthetase genes and led to consequent accumulation of asparagine, a nitrogen-rich amino acid is known to facilitate the reallocation of nitrogen resources from senescent to young growing organs. Hypothesizing these combined host responses to restrain foreign protein accumulation, an attempt was made to support nitrogen reduction in CoVLP-producing leaves by co-expressing a constitutively active, light-insensitive form of the nitrate reductase. We show this strategy to increase S protein accumulation in leaf tissues, thereby suggesting that boosting nitrogen metabolism of agroinfiltrated leaves improves recombinant protein yields in *N. benthamiana*.

## Introduction

Senescence, as the final stage of leaf development, is a programmed form of plant cell death (PCD) characterized by the orderly disassembly of cellular components (Woo *et al*., 2019; Guo *et al*., 2021). Before termination, plant cells operate a major metabolic shift, from anabolism to catabolism, associated with chlorophyll degradation and the breakdown of macromolecules including lipids, proteins, and nucleic acids. These catalytic processes, which result in a progressive decline of leaf photosynthetic capacity, allow on the other hand for the recycling of primary metabolic resources and their reallocation to non-senescent organs. From a development standpoint, leaf senescence not only represents a degenerative process, but also a strategy to support plant fitness through the efficient reuse of key nutrients in growing tissues, including nitrogen (N).

As a constituent of amino acids, proteins, nucleic acids, and chlorophyll, N is a central element for plant development. Highly abundant in the lithosphere and the atmosphere, this element is however mainly present in inorganic forms not directly usable by plants for metabolic purposes (Mengel *et al*., 2001). Inorganic N is first absorbed by the root system as ammonium (NH_4_^+^) or nitrate (NO_3_^−^), the latter form often being the most abundant in the soil and mainly originating from bacterial nitrification and soil fertilization (Good *et al*., 2004). In root cells, NO_3_^−^ is reduced to produce amino acids that are either used directly for protein biosynthesis or translocated to the shoot following uploading in the xylem. Alternatively, absorbed NO_3_^−^ anions can be directly uploaded in the xylem for immediate transport and later reduction in shoot tissues (Tegeder, 2014). To be incorporated into amino acids, NO_3_^−^ has to be first reduced to NH_4_^+^ (Zayed *et al*., 2023), a compartmentalized process that starts in the cytosol where nitrate reductase (NR) reduces NO_3_^−^ to nitrite (NO_2_^−^). Nitrite ions then translocate into the plastids (or the chloroplasts) for further reduction to NH_4_^+^ by nitrite reductase (NiR), a prerequisite for subsequent assimilation in the amino acid metabolism.

Under a reduced state compatible with cellular metabolism, NH_4_^+^ is conjugated to glutamate by glutamine synthetase (GLN), a reaction producing glutamine. Combined to 2-oxoglutarate, glutamine then generates two molecules of glutamate in a reaction catalyzed by glutamate synthase (GLU), also termed glutamine 2-oxoglutarate aminotransferase (GOGAT). Aside from the so-called GLN/GLU (GOGAT) cycle, N assimilation is also influenced by asparagine synthetase (ASN), which performs the ATP-dependent transfer of the amine group of glutamine on aspartate to produce glutamate and asparagine (Lomelino *et al*., 2017). Used directly for anabolic functions or exported to alternative tissues to sustain growth, amino acids also serve as building blocks for the storage of N resources in the polypeptide backbone of highly abundant proteins such as ribulose-1,5-bisphosphate carboxylase/oxygenase (RuBisCO) or chloroplastic GLN (Tegeder and Masclaux-Daubresse, 2018).

During senescence, free amino acids are released from the breakdown of cellular proteins, an activity mediated by the hydrolytic action of proteolytic enzymes such as C1A cysteine proteases (CPs) (Roberts *et al*., 2012). Amino acids are then reallocated to alternative tissues for reassimilation in the primary metabolism (Havé *et al*., 2017; Tegeder and Masclaux-Daubresse, 2018). In ageing leaves, total amino acid content progressively decreases in the leaf blade, while increasing in the phloem sap. Metabolomic analyses suggest that N reallocation in several plant species occurs preferentially through the translocation of specific amino acids in the phloem, subsequent to an extensive modulation of the overall amino acid spectrum in senescing cells. For instance, glutamate and aspartate contents rise at the onset of leaf senescence in *Arabidopsis thaliana*, before decreasing at later stages (Diaz *et al*., 2005; Watanabe *et al*., 2013). Conversely, glutamine and asparagine contents steadily increase, indicative of an active interconversion of glutamate to glutamine and of aspartate to asparagine in senescing tissues. Harbouring two N atoms and being among the most abundant amino acids in the phloem, glutamine and asparagine play a central role in the reallocation of N resources to younger organs during senescence (Havé *et al*., 2017).

As a fine-tuned and multi-faceted process, leaf senescence is both the result of a predetermined genetic program and a response to external factors such as biotic and abiotic stress stimuli that favor or even expedite the ageing process (Schippers *et al*., 2015; Woo *et al*., 2019). This was recently illustrated with the transient protein expression host *Nicotiana benthamiana* agroinfiltrated to produce influenza virus-like particles (VLPs) as vaccine candidate (Hamel *et al*., 2024). The agroinfiltration procedure, which relies on leaf tissue infiltration with the bacterial gene vector *Rhizobium radiobacter* (commonly known and hereafter referred to as *Agrobacterium tumefaciens*), induces a strong defense response associated with the upregulation of oxylipin-related genes and signalling crosstalk between key stress hormone pathways. The production of influenza VLPs, and to some extent the agroinfiltration procedure *per se*, were also suggested to induce senescence-related genes in young, growing leaves of *N. benthamiana* (Hamel *et al*., 2024). These included ethylene biosynthesis genes, a homolog of Arabidopsis *ORESARA 1* (*AtORE1*) known to antagonize senescence-repressing effects of other transcription factors (TFs) (Rauf *et al*., 2013), a homolog of Arabidopsis *STAYGREEN 1* (*AtSGR1*) involved in chlorophyll degradation (Sakuraba *et al*., 2015), and close homologs of *Nicotiana tabacum* (tobacco) *ASN* genes involved in the remobilization of N resources towards non-senescent organs (Bovet *et al*., 2019). From a practical standpoint, the upregulation of senescence-related genes during foreign protein expression raises questions about the net impact of stress-induced leaf senescence on recombinant protein yields in plants used as protein ‘biofactories’. In particular, the protein degradation-promoting effect of this process and the possible negative impact of N remobilization to alternative organs on the availability of free amino acids for *de novo* protein biosynthesis underline the relevance of a more detailed account on this process, as a prerequisite for the eventual implementation of mitigation strategies in a molecular farming context.

Soon after the beginning of the coronavirus disease 2019 (COVID-19) pandemic, the biopharmaceutical company Medicago developed a protective vaccine known as Covifenz^®^ (Hager *et al*., 2022). This first plant-made vaccine authorized for use in humans relied on *N. benthamiana* leaves agroinfiltrated to transiently express the Spike (S) protein of Severe Acute Respiratory Syndrome Coronavirus 2 (SARS-CoV-2). The process also relied on co-expression of the viral suppressor of RNA silencing P19 (Silhavy *et al*., 2002), which prevented silencing of the *S* transgene delivered by *Agrobacterium*. Upon expression, the secreted S protein results in spontaneous formation of the so-called coronavirus-like particles (CoVLPs), which bud from the plasma membrane (PM) of plant cells. Using RNAseq and time course leaf sampling over several days, we recently characterized the molecular responses of *N. benthamiana* leaf cells agroinfiltrated to express P19 only, or P19 in combination with recombinant S protein (see accompanying manuscript; Hamel *et al*., 2025). This work revealed CoVLP production to deeply affect the physiological status of plant cells, notably through a strong activation of their immune system.

Further exploiting our leaf transcriptome datasets, we here show that CoVLP production also stimulates leaf senescence through the induction of TF genes involved in senescence, the upregulation of senescence-associated genes (SAGs) with varied functions, and the biosynthesis of senescence-related proteases. Expression of the S protein also results in the downregulation of primary metabolism-associated genes involved in photosynthesis, N reduction, and N assimilation, concomitant with the upregulation of senescence-related *ASN* genes and consequent accumulation of asparagine in agroinfiltrated leaf tissues. Expecting these combined responses in the plant to divert N and amino acid resources away from protein biosynthesis, including the recombinant S protein to be produced, we overexpressed a constitutively active, light-insensitive version of an *N. benthamiana* NR that was otherwise downregulated upon CoVLP accumulation. This approach aimed at mitigating the downregulating effect of stress-induced senescence on NO_3_^−^ reduction, a key limiting step in the N assimilation pathway. Co-expression experiments confirmed the deregulated NR to significantly increase S protein accumulation in leaf tissues, unlike the wild-type, dark-sensitive enzyme showing no effect. Our results thus suggest senescence-related responses and constrained N assimilation to indeed limit foreign protein accumulation in *N. benthamiana* leaves, an effect that can genetically be rescued by supporting N reduction in agroinfiltrated tissues.

## Results

In an associated study, *N. benthamiana* plants agroinfiltrated to transiently co-express P19 and the S protein were sampled to characterize molecular responses induced by CoVLP production in leaves (CoVLP samples; Hamel *et al*., 2025). Based on systematic numbering of primary (P) leaves and distribution of the S protein within the canopy, leaves P9 and P10 were selectively harvested in a time course fashion at 2, 3, 4, 5, and 6 days post-infiltration (DPI). For each time point, P9 and P10 leaves of agroinfiltrated plants solely expressing P19 were also collected as controls (P19 samples). As a reference point, P9 and P10 leaves of non-infiltrated (NI) plants harvested prior to infiltration were also analysed (NI samples at 0 DPI). After evaluating leaf symptoms and formally confirming expression of both the transgenes and recombinant proteins, RNAseq analyses and real time quantitative polymerase chain reaction (RTqPCR) measurements were performed to profile plant gene expression throughout the course of foreign protein accumulation. A thorough assessment of the resulting leaf transcriptomes confirmed the differential regulation of several cellular processes in the host plant, including the activation of immune responses and the onset of signalling crosstalk between important stress hormone pathways (Hamel *et al*., 2025).

### Upregulation of senescence-associated TF genes

Our RNAseq datasets revealed the induction of several TF genes upon S protein expression, including WRKY family members associated to plant immunity (Hamel *et al*., 2025). Also flagged as upregulated were genes belonging to the *NO APICAL MERISTEM* (*NAM*), *ARABIDOPSIS THALIANA ACTIVATION FACTOR 1* (*ATAF1*), and *CUP SHAPED COTYLEDON 2* (*CUP2*) family. Simply referred to as *NACs*, these genes form a large family of TFs in plants, which for instance includes more than 100 members in Arabidopsis (Ooka *et al*., 2003). NACs are involved in a variety of cellular processes, including the promotion of leaf senescence in Arabidopsis and tobacco (Podzimska-Sroka *et al*., 2015; Li *et al*., 2018). Here, several *NACs* were induced upon S protein expression (Table S1), including the closely related genes *NbNAC87* (Niben101Scf02762g01001) and *NbNAC81* (Niben101Scf15435g00012), encoding homologs of senescence-promoting NACs in other plant species. Named after *AtNAC87* (At5g18270) and *AtNAC81* (At5g08790), their closest respective homolog in Arabidopsis, *NbNAC87* and *NbNAC81* were upregulated specifically in CoVLP samples to reach maximum transcript levels at 3 or 4 DPI (Figure 1a). We recently identified ‘early’ and ‘late’ TF genes in *N benthamiana* leaves expressing the S protein (Hamel *et al*., 2025). Based on their expression profiles, *NbNAC87* and *NbNAC81* were considered as early TFs in the current expression system. In Arabidopsis, AtNAC87 works as a transcriptional activator to promote age-related leaf senescence (Chen *et al*., 2023). As for AtNAC81, transient reporter gene assays showed this TF to act as a transcriptional activator or repressor, depending on its target gene promoter (Nagahage *et al*., 2018). Also known as ARABIDOPSIS THALIANA ACTIVATION FACTOR 2 (ATAF2), AtNAC81 was shown to promote age-related and dark-induced leaf senescence in Arabidopsis (Nagahage *et al*., 2020).

**Figure 1.**
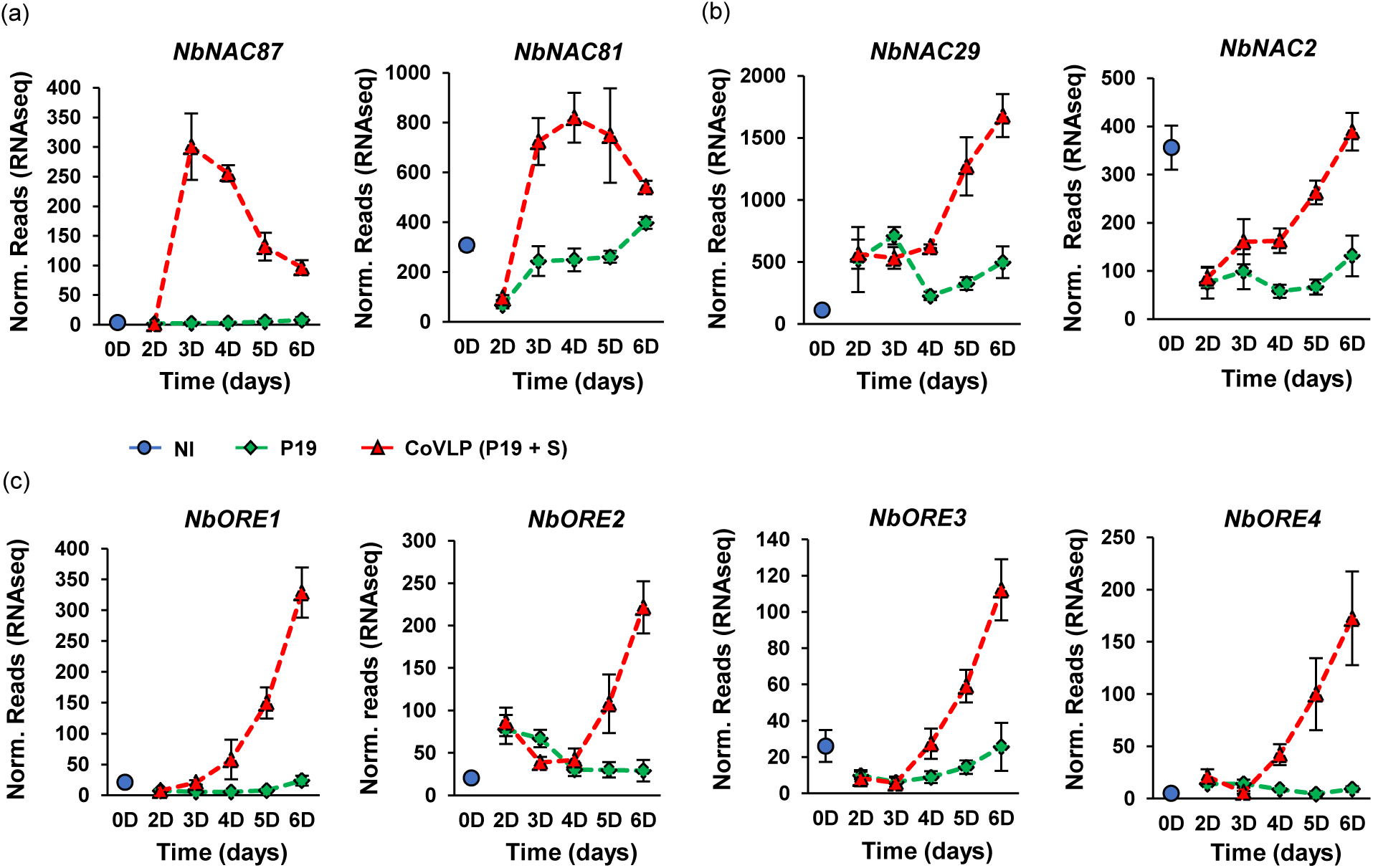
Upregulation of senescence-related *NACs*. Expression profiles of TF genes encoding early (a) or late (b) NACs. Expression profiles of late *NACs* encoding TFs closely related to Arabidopsis AtORE1 and AtORS1 are also shown (c). For each time point in days (D) post-infiltration, RNAseq results are expressed in normalized (norm.) numbers of reads ± sd. Conditions are as follows: NI: non-infiltrated samples (blue); P19: agroinfiltrated samples expressing P19 only (green); CoVLP: agroinfiltrated samples co-expressing P19 and recombinant S protein (red).

Our RNAseq data also revealed the upregulation of late *NACs*, with transcript levels still on the rise at the end of the sampling period. These included *NbNAC29* (Niben101Scf10322g01012), a gene induced more strongly in CoVLP samples than in P19 samples (Figure 1b). In Arabidopsis, the closest homolog of *NbNAC29* is *AtNAC29* (At1g69490), a senescence-promoting gene also known as *ARABIDOPSIS NAC-LIKE ACTIVATED BY APETALA3/PISTILLATA* (*AtNAP*) (Guo and Gan, 2006). Similarly, functional analyses revealed senescence-promoting functions for *NaNAC29* (Ma *et al*., 2021) and *NtNAC80* (Li *et al*., 2018), the closest homologs of *NbNAC29* in *Nicotiana attenuata* and tobacco, respectively. Also categorized as a late *NAC*, *NbNAC2* (Niben101Scf00799g00011) was quickly downregulated upon agroinfiltration in both P19 and CoVLP samples (Figure 1b). Concomitant with progressive accumulation of the S protein in leaf tissues (Hamel *et al*., 2025), this gene was however reinduced in CoVLP samples. The closest homolog of this gene in Arabidopsis, referred to as *ATAF1* or *AtNAC2* (At1g01720), encodes another key promoter of the leaf senescence process (Garapati *et al*., 2015).

Four highly homologous *NACs* termed *NbORE1* (Niben101Scf01277g00002), *NbORE2* (Niben101Scf04133g02029), *NbORE3* (Niben101Scf00063g07011), and *NbORE4* (Niben101Scf05060g08015) were also induced at late time points, a response essentially specific to CoVLP samples (Figure 1c). The products of these genes are closely related to the well-known inducers of leaf senescence in Arabidopsis, AtORE1 (At5g39610) (Kim *et al*., 2009; Balazadeh *et al*., 2010) and *ORESARA 1 SISTER 1* (*AtORS1*) (At3g29035; Balazadeh *et al*., 2011). The upregulation of these genes further links S protein expression to the activation of a complex regulatory network that leads to senescence in CoVLP-producing leaves.

### Upregulation of SAGs and other senescence-related genes

When reaching the senescence stage, leaf cells undergo substantial changes in their gene expression profiles (Woo *et al*., 2019). Whereas most genes normally expressed in non-senescent leaves become downregulated, a subset of genes is on the opposite induced to coordinate the senescence program. Several transcriptomics studies have been conducted to identify these regulatory genes, often referred to as SAGs (Gepstein *et al*., 2003; Guo *et al*., 2004; Buchanan-Wollaston *et al*., 2005; Breeze *et al*., 2011). The Leaf Senescence Database (Liu *et al*., 2011), as an integrated and searchable gene database, for instance referenced over 31,000 putative SAGs distributed across ∼150 plant species in its 2023 update (https://ngdc.cncb.ac.cn/lsd/).

RNAseq datasets here showed many SAG homologs to be upregulated in CoVLP samples (Table S1), including *NbSAG14a* (Niben101Scf02838g07002) and *NbSAG14b* (Niben101Scf10688g02001; Figure 2a). Closely related to each other, these genes are also close homologs of *AtSAG14* (At5g20230), which encodes an endomembrane protein positively regulating dark-induced leaf senescence in Arabidopsis (Hao *et al*., 2022). Also identified were *NbSAG15a* (Niben101Scf07623g01034) and *NbSAG15b* (Niben101Scf02581g05002), both downregulated shortly after agroinfiltration but later reinduced in CoVLP samples (Figure 2a). Closely related to each other, sequence analysis revealed *NbSAG15a* and *NbSAG15b* to be close homologs of *AtSAG15* (At5g51070), an Arabidopsis gene also known as *EARLY RESPONSIVE TO DEHYDRATION 1* (*AtERD1*). Located in the stroma, AtERD1 works as a molecular chaperone for a protease complex involved in the degradation of abundant chloroplastic proteins during senescence (Nakashima *et al*., 1997). Upon CoVLP accumulation, induced SAGs also included *NbSAG20* (Niben101Scf00773g08003) and *NbSAG201* (Niben101Scf09665g00001; Figure 2a), named after their closest Arabidopsis homologs *AtSAG20* (At3g10985) and *AtSAG201* (At2g45210), respectively. *AtSAG20* encodes a wound-induced protein shown to be strongly upregulated during leaf senescence (Breeze *et al*., 2011). *AtSAG201,* also termed *AtSAUR36*, belongs to the *small auxin upregulated RNA* (*SAUR*) gene family and was reported to promote auxin-dependent leaf senescence in Arabidopsis (Hou *et al*., 2013).

**Figure 2.**
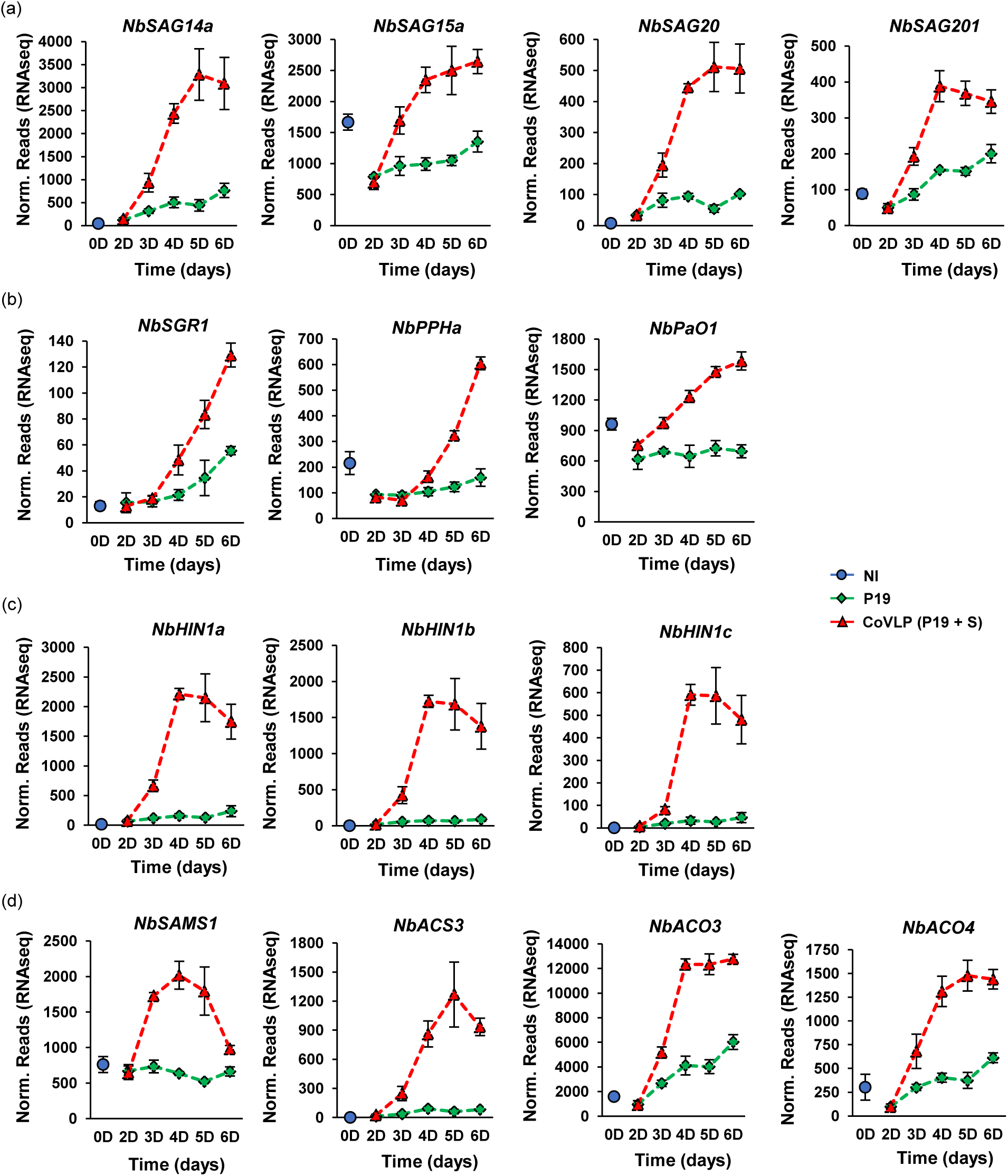
Upregulation of senescence-related genes. (a) Expression profiles of SAGs with varied functions. Expression profiles of genes involved in senescence-related degradation of the chlorophyll (b), senescence-related PCD (c), and ethylene biosynthesis (d) are also shown. For each time point in days (D) post-infiltration, RNAseq results are expressed in normalized (norm.) numbers of reads ± sd. Conditions are as follows: NI: non-infiltrated samples (blue); P19: agroinfiltrated samples expressing P19 only (green); CoVLP: agroinfiltrated samples co-expressing P19 and recombinant S protein (red).

In addition to the genes formally labelled as SAGs, leaf senescence induces many other regulatory genes, several of them implicated in chlorophyll degradation, senescence-associated PCD, or ethylene biosynthesis (Woo *et al*., 2019; Guo *et al*., 2021). Accordingly, RNAseq data here revealed several homologs of these genes to be strongly induced upon CoVLP production (Table S1), including homologs of Arabidopsis genes *AtSGR1* (At4g22920), *PHEOPHYTINASE* (*AtPPH*) (At5g13800), and *PHEOPHORBIDE a OXYGENASE* (*AtPaO*) (At3g44880), which encode chlorophyll catabolic enzymes involved in the degreening of leaf tissues during senescence (Pružinská *et al*., 2003; Schelbert *et al*., 2009; Sakuraba *et al*., 2015). Compared to NI samples at 0 DPI, the *AtSGR1* homolog *NbSGR1* (Niben101Scf00490g01040) was induced in both P19 and CoVLP samples, but with upregulation levels much higher in the latter case (Figure 2b). By comparison, *AtPPH* homolog *NbPPHa* (Niben101Scf12801g00006) and *AtPaO* homolog *NbPaO1* (Niben101Scf00436g10019) were slightly repressed shortly after agroinfiltration in both P19 and CoVLP samples, but progressively reinduced in CoVLP samples (Figure 2b) as the S protein started to accumulate (Hamel *et al*., 2025).

RNAseq also revealed the strong upregulation of three closely related genes, *NbHIN1a* (Niben101Scf08020g06001), *NbHIN1b* (Niben101Scf04717g02003), and *NbHIN1c* (Niben101Scf08020g05009) (Figure 2c). Specific to CoVLP samples, these genes encode hydroxyproline-rich glycoproteins with high identity to the gene product of tobacco *HARPIN-INDUCED 1* (*NtHIN1*), a PCD-associated gene highly active in senescent leaves and flowers (Takahashi *et al*., 2004).

Several genes involved in ethylene biosynthesis were induced upon CoVLP production (Table S1), including the *S-adenosylmethionine synthetase* (*SAMS*) gene *NbSAMS1* (Niben101Scf02502g04001) and the *1-aminocyclopropane-1-carboxylic acid synthase* (*ACS*) gene *NbACS3* (Niben101Scf09512g03008) (Figure 2d). *ACC oxidase* (*ACO*) genes were also upregulated at much higher levels in CoVLP samples compared to P19 samples (Table S1), including *NbACO3* (Niben101Scf02543g01008) and *NbACO4* (Niben101Scf08039g01005; Figure 2d). During the final stages of leaf senescence, ethylene production is known to increase along with the expression of ethylene biosynthesis genes (Van der Graaff *et al*., 2006; Breeze *et al*., 2011). Taken together, our transcriptomics results showed that senescence-related genes of varied functions were upregulated during foreign protein expression, especially when the S protein was expressed.

### Upregulation of CP genes and increased CP abundance and activity in leaves

Proteolysis allows for the recycling of N resources during leaf senescence, in addition of a regulatory role in senescence-associated PCD (Roberts *et al*., 2012). At the cellular level, increased protease levels and activities are typically preceded by the upregulation of their corresponding genes, with CP-encoding genes consistently representing the most abundant class of upregulated protease genes during senescence (Guo *et al*., 2004; Havé *et al*., 2017). In Arabidopsis, senescence-associated CP genes include *RESPONSIVE TO DEHYDRATION 19a* (*AtRD19a*; At4g39090), and its close homologs *AtRD19b* (At2g21430) and *AtRD19c* (At4g16190). Also induced is the gene *RESPONSIVE TO DEHYDRATION 21a* (*AtRD21a*; At1g47128), and its close homologs *AtRD21b* (At5g43060) and *AtRD21c* (At3g19390) (Breeze *et al*., 2011; Pružinská *et al*., 2017). Homologs of Arabidopsis *RD19* and *RD21* genes have also been reported to be involved in the senescence of other plant species, including in *Brassica oleracea* (broccoli) following floret harvesting (Coupe *et al*., 2003). Accordingly, RNAseq data here revealed the preferential upregulation in CoVLP samples of protease genes encoding close homologs of RD19-like proteases (Table S1; Figure S1a), namely *NbRD19a* (Niben101Scf00973g01002), *NbRD19b* (Niben101Scf01701g00013), and *NbRD19c* (Niben101Scf06228g02005) (Figure 3a). Also termed *NbCYP2*, *NbRD19b* was previously involved in responses to pathogen infection in *N. benthamiana* (Hao *et al*., 2006). Also preferentially induced in CoVLP samples were three CP genes termed *NbRD21a* (Niben101Scf04007g01012), *NbRD21b* (Niben101Scf12813g00004), and *NbRD21c* (Niben101Scf09885g00008) (Table S1; Figure 3b), all encoding products closely related to RD21-like proteases (Figure S1a).

**Figure 3.**
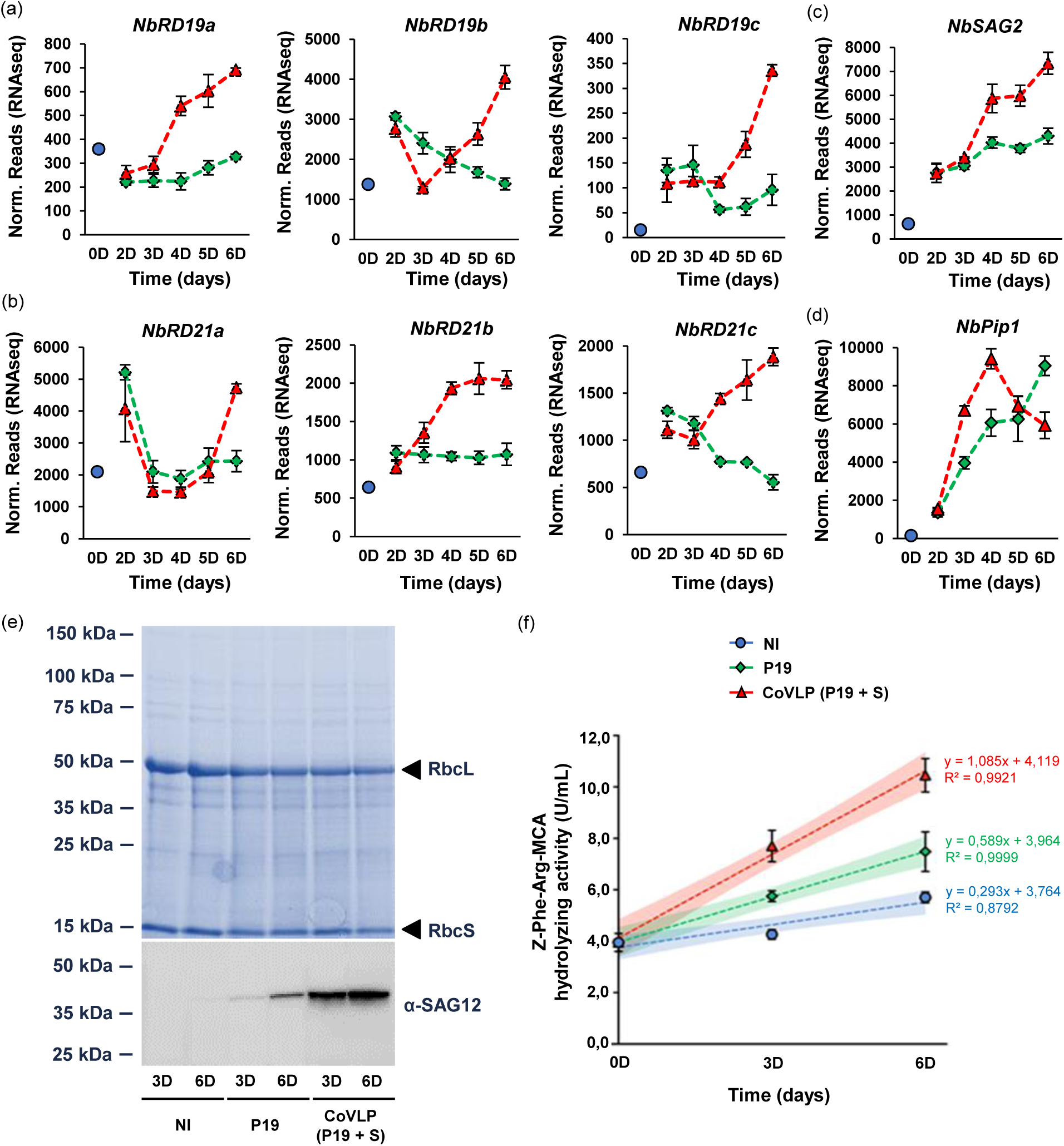
Upregulation of CP genes, accumulation of SAG12-like proteases, and increase of CP activity. Expression profiles of CP genes encoding RD19-(a), RD21-(b), SAG2-(c), and SAG12-like (d) homologs, all of which are involved in senescence-related proteolysis. For each time point in days (D) post-infiltration, RNAseq results are expressed in normalized (norm.) numbers of reads ± sd. (e) Total protein extracts following SDS-PAGE and Coomassie blue-staining (upper panel). Arrows highlight RuBisCO small (RbcS) and large (RbcL) subunits. The western blot (lower panel) depicts accumulation of SAG12-like proteases, which breakdown RuBisCO during senescence. (f) CP activity as measured by hydrolysis of the synthetic fluorogenic substrate *Z*-Phe-Arg-MCA and standard curves made using recombinant human cathepsin L. Results are expressed in enzymatic activity units (U) per mL of total protein extract. For each condition, the dashed line corresponds to the simple linear regression of the data at each time point. Colored areas delimit 95% confidence intervals of the linear regressions. First degree equations and deduced coefficient of determination (R^2^) are shown on the right. Conditions are as follows: NI: non-infiltrated samples (blue); P19: agroinfiltrated samples expressing P19 only (green); CoVLP: agroinfiltrated samples co-expressing P19 and recombinant S protein (red).

In Arabidopsis, *SENESCENCE-ASSOCIATED GENE 2* (*AtSAG2*; At5g60360) is also known to be strongly induced during leaf senescence (Gepstein *et al*., 2003; Breeze *et al*., 2011; Pružinská *et al*., 2017). The encoded protein, also referred to as ALEURAIN-LIKE PROTEASE (AtALP), actually accounts for most of the CP activity in senescent leaves of Arabidopsis, along with AtRD21a (Pružinská *et al*., 2017). In *N. benthamiana*, the *NbSAG2* (Niben101Scf01445g00016) gene product, also termed NbCYP1 (Hao *et al*., 2006), was here identified as the closest homolog of AtSAG2 (Figure S1a). Compared to NI samples at 0 DPI, *NbSAG2* was similarly induced in both P19 and CoVLP samples up to 3 DPI (Figure 3c). At later time points, the expression of this gene continued to increase in P19 samples, but again at much lower levels compared to CoVLP samples.

In *Oryza sativa* (rice), *SENESCENCE-ASSOCIATED GENE 39* (*OsSAG39*; Os04g12660) encodes a CP specifically expressed during leaf senescence (Liu *et al*., 2010). OsSAG39 shares ∼55% homolog with Arabidopsis *SENESCENCE-ASSOCIATED GENE 12* (*AtSAG12*; At5g45890), which is also specifically induced during leaf senescence (Breeze *et al*., 2011; Pružinská *et al*., 2017). Widely employed as a senescence marker, AtSAG12 localizes in senescence-associated vacuoles involved in the degradation of highly abundant stromal proteins, including RuBisCO (Otegui *et al*., 2005; James *et al*., 2018). In *N. benthamiana*, several CPs cluster with SAG12-like proteases (Grosse-Holz *et al*., 2018), including the product of *Phytophthora-inhibited protease 1* (*NbPip1*; Niben101Scf08921g02024) (Figure S1a), a gene involved in activation of the hypersensitive response (HR) (Xu *et al*., 2012). Compared to NI samples at 0 DPI, *NbPip1* was here induced in both P19 and CoVLP samples, with maximum expression reaching similar levels in both conditions, albeit two days earlier in CoVLP samples (Figure 3d).

Among the numerous *SAG12-like* genes *in N. benthamiana*, *NbSAG12* (Niben101Scf10219g00026) encodes a CP most closely related to AtSAG12 (Figure S1a). RNAseq and RTqPCR analyses showed the expression of *NbSAG12* to remain barely detectable under our experimental conditions, including in CoVLP samples at 6 DPI. We assumed this to be likely explained by the selective sampling of young and actively growing leaves for the transcriptomics analyses (Hamel *et al*., 2025). To further assess whether SAG12-like proteases are part of the cellular responses to foreign protein expression, a western blot was performed using antibodies specific to conserved regions of SAG12-like proteases. To favor signal detection, protein extracts were produced using pooled leaf biomass from whole plant shoots, so older leaves were also included in the tested material. Sampling was performed on P19-and CoVLP-expressing plants harvested at 3 and 6 DPI, using NI plants of the same age as controls. Immunoblotting revealed higher accumulation of SAG12-like proteases in CoVLP samples compared to P19 samples, a result respectively observed at both sampling points (lower panel of Figure 3e). For NI samples, no signal was detected at 3 DPI, and a barely detectable signal was observed at 6 DPI (Figure 3e).

In Arabidopsis, functional analyses also revealed three *cathepsin B* (*CathB*) genes termed *AtCathB1* (At1g02300), *AtCathB2* (At1g02305), and *AtCathB3* (At4g01610) to redundantly promote different forms of programmed PCD, including developmentally regulated PCD forms associated with leaf senescence (McLellan *et al*., 2009; Ge *et al*., 2016; Cai *et al*., 2017). CathB inhibitors and virus-induced gene silencing also revealed that CathBs could trigger HR following pathogen infection in *N. benthamiana* (Gilroy *et al*., 2007). Here, RNAseq data highlighted three CP genes (Table S1) whose encoded products cluster with Arabidopsis CathBs (Figure S1a) and bear the occluding loop typical of CathB family members (Figure S1b) (Coppola *et al*., 2024). Whereas *NbCathB1* (Niben101Scf02976g00007) and *NbCathB3* (Niben101Scf03867g02046) were essentially repressed during recombinant protein expression, *NbCathB2* (Niben101Scf01455g01021) was induced in both P19 and CoVLP samples, again at much higher levels upon S protein expression (Figure S1c).

To confirm that the upregulation of CP genes resulted in increased abundance of the encoded proteases during foreign protein expression, we searched proteomics datasets recently generated using isobaric tags for relative and absolute quantitation (iTRAQ) labelling (Hamel *et al*., 2025). Using NI samples at 0 DPI as a control, pairwise comparisons were performed for P19 and CoVLP samples at 6 DPI. Pairwise comparisons between P19 and CoVLP samples at 6 DPI were also performed to formally compare relative induction rates in the two sample types. Differentially expressed proteins were defined as all candidates whose differential expression adjusted p-value (padj) was below 0.05, corresponding to a false discovery rate below 5%. CPs showing increased accumulation were then extracted from the resulting lists of regulated proteins. Overall, seven CPs could be identified by this approach (Table S2), including NbPip1 showing the highest increase in accumulation in both P19 and CoVLP samples at 6 DPI. Interestingly, NbPip1 accumulation levels were similar in both conditions (Table S2), in line with similar maximum levels of *NbPip1* transcripts assayed in the corresponding samples (Figure 3d). By contrast, NbCathB2, NbSAG2, NbRD21b, and NbRD21c accumulated at significantly higher levels in CoVLP samples than in P19 samples at 6 DPI (Table S2). Considering the expression profiles of corresponding genes (Table S1; Figure 3b,c; Figure S1c), our transcriptomics and proteomics data correlated extensively, further confirming that CoVLP production in leaves not only resulted in the upregulation of CP genes but also in the enhanced accumulation of their encoded proteases.

To further demonstrate the importance of CPs during foreign protein expression, changes in CP (cathepsin L-like) activity were monitored using the synthetic fluorogenic peptide *Z*-Phe-Arg 7-amido-4-methylcoumarin (*Z*-Phe-Arg-MCA) as a substrate. As for the immunodetection of SAG12-like proteases (Figure 3e), CP activity was measured at 3 and 6 DPI using pooled leaf biomass from whole plant shoots expressing P19 only, or P19 along with the S protein. We also analysed the extracts of NI plants harvested prior to infiltration, as well as age-related controls consisting of NI plants harvested 3 and 6 days after infiltration of the P19 and CoVLP plants. Cathepsin L activity in NI plants remained mostly unchanged when comparing samples harvested at 0 and 3 DPI (Figure 3f). By contrast, cathepsin L activity had significantly increased in NI plants harvested at 6 DPI, suggesting plant ageing to result in increased CP activity regardless of the agroinfiltration procedure. For P19 and CoVLP plants, cathepsin L activity also increased with time, but the activity levels were significantly higher than in the NI plants at the corresponding time points. At 3 DPI and even more at 6 DPI, direct comparison between P19 and CoVLP plants also revealed that CP activity was significantly higher in plants expressing the S protein (Figure 3f). Together, our data on the expression of CP genes, the accumulation of SAG12-like proteases, the relative abundance of CPs, and the increased CP activity in plant extracts clearly indicated an upregulated effect of foreign protein expression on CP accumulation and activity in the host plant, especially during CoVLP accumulation. While increased CP abundance and activity may be linked to the activation of plant immunity after the agroinfiltration (Hamel *et al*., 2025), these responses could also be seen as a manifestation of the leaf senescence process (Guo *et al*., 2004; Havé *et al*., 2017).

### Downregulation of chloroplast-related genes

Functional analyses in Arabidopsis revealed that GOLDEN2-LIKE (GLK) TFs AtGLK1 (At2g20570) and AtGLK2 (At5g44190) regulate chloroplast development and maintenance of the photosynthetic functions (Waters *et al*., 2009; Rauf *et al*., 2013). For instance, *atglk1/atglk2* double mutants are pale green and present a deficient photosynthetic apparatus, while the overexpression of *GLKs* increases the expression of photosynthesis-associated genes (PAGs) and delays leaf senescence. We recently identified four *GLK* genes in *N. benthamiana*, *NbGLK1a* (Niben101Scf11137g00006), *NbGLK1b* (Niben101Scf09774g03004), *NbGLK2a*

(Niben101Scf06721g00011), and *NbGLK2b* (Niben101Scf01462g01010), named after their closest homolog in Arabidopsis (Hamel *et al*., 2024). Our RNAseq datasets here revealed all four *GLK* genes to be differentially expressed at various time points during foreign protein expression (Table S1). At 2 DPI, *GLKs* were all induced in both P19 and CoVLP samples compared to NI samples at 0 DPI (Figure 4a). Given the absence of S protein at 2 DPI (Hamel *et al*., 2025), we hypothesized these early responses to result from a mechanical stress caused by agroinfiltration, or from perception of the *Agrobacterium* vector by plant cells. As the expression phase progressed, *GLK* gene expression gradually decreased, a response generally more pronounced in CoVLP samples than in P19 samples (Figure 4a). For *GLKs* significantly expressed in NI plants at 0 DPI (*NbGLK1a*, *NbGLK1b*, and *NbGLK2a*), the later decrease in gene expression led to transcript levels lower than those measured in NI plants prior to infiltration, especially when the S protein was expressed (Figure 4a).

**Figure 4.**
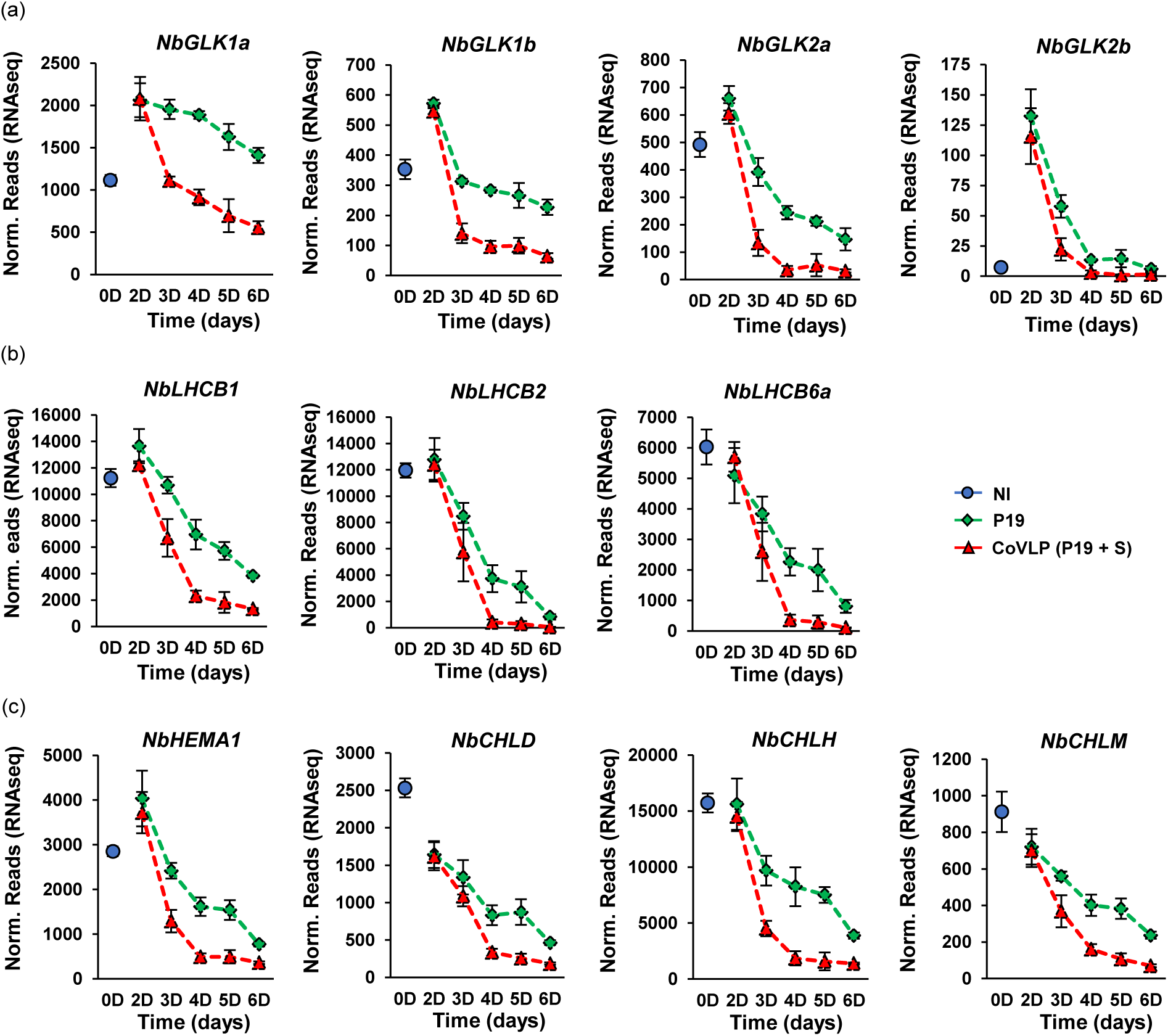
Downregulation of chloroplast-related genes. Expression profiles of genes encoding TFs of the GLK family (a). Expression profiles of PAGs are also shown, namely those encoding chloroplastic proteins involved in light capture (b), or chlorophyll biosynthesis (c). For each time point in days (D) post-infiltration, RNAseq results are expressed in normalized (norm.) numbers of reads ± sd. Conditions are as follows: NI: non-infiltrated samples (blue); P19: agroinfiltrated samples expressing P19 only (green); CoVLP: agroinfiltrated samples co-expressing P19 and recombinant S protein (red).

To further examine the regulation of chloroplast-related genes, we profiled the expression of PAGs involved in light capture or biosynthesis of the chlorophyll pigments (Table S1). Some of the examined genes actually corresponded to the homologs of direct genetic targets of AtGLK1 (Waters *et al*., 2009). As depicted for the *LIGHT-HARVESTING CHLOROPHYLL a/b BINDING* (*LHCB*) genes *NbLHCB1* (Niben101Scf08088g04018), *NbLHCB2* (Niben101Ctg16374g00003), and *NbLHCB6a* (Niben101Scf02513g08007), RNA transcripts were rapidly depleted during foreign protein expression, especially in CoVLP samples (Figure 4b). For genes involved in chlorophyll biosynthesis, examined candidates included *NbHEMA1* (Niben101Scf03068g00024) encoding a glutamyl-tRNA reductase, *NbCHLD* (Niben101Scf01209g04002) and *NbCHLH* (Niben101Scf04388g00011) encoding subunits of the magnesium chelatase protein complex, as well as *NbCHLM* (Niben101Scf03548g02031) encoding a magnesium-protoporphyrin IX methyltransferase. In all cases, the expression profiles revealed severe transcript depletion upon foreign protein expression, with again the stronger effects seen upon S protein expression (Figure 4c). Overall, these observations on chloroplast-related genes suggested recombinant protein expression to result in an alteration of the photosynthetic functions in chloroplasts, especially upon CoVLP accumulation. While altered chloroplast functions could result from a trade-off relationship linked to the activation of plant immunity (Hamel *et al*., 2025), they could also represent, similar to the upregulation of CP genes above, a key hallmark of the leaf senescence process (Woo *et al*., 2019; Guo *et al*., 2021).

### Upregulation of ASN genes involved in N remobilization

During senescence, asparagine plays a central role in the remobilization of N resources to metabolically active organs (Figure 5a) (Havé *et al*., 2017). In Arabidopsis (Lin and Wu, 2004; Diaz *et al*., 2005; Watanabe *et al*., 2013), *Helianthus annuus* (sunflower) (Moschen *et al*., 2016), and tobacco (Bovet *et al*., 2019), this amino acid accumulates in ageing leaves or as a result of environmental conditions that trigger leaf senescence. Since asparagine biosynthesis relies on the catalytic activity of ASN (Lomelino *et al*., 2017), the expression profiles of *ASN* genes in leaves were investigated. An analysis of the *N. benthamiana* genome revealed this gene family to comprise six members, matching the six *ASN* genes of tobacco (Bovet *et al*., 2019). A Phylogenetic analysis showed *N. benthamiana* ASNs to segregate in two clusters, Cluster I and Cluster II, similar to the corresponding segregation pattern of their ASN homologs in Arabidopsis (Gaufichon *et al*., 2010) and other Solanaceae (Figure 5b). A close assessment of our RNAseq data revealed contrasted outcomes for the expression profiles of *ASN* gene members from each cluster (Table S1). Whereas *NbASN1* (Niben101Scf00573g01003), *NbASN2* (Niben101Scf11860g00013), and *NbASN4* (Niben101Scf05880g03001) from Cluster I exhibited no or very weak expression in NI leaves at 0 DPI (Figure 5c), *NbASN5* (Niben101Scf01936g07002) and *NbASN6* (Niben101Scf02268g00002) from Cluster II showed higher basal expression levels in these leaf tissues (Figure 5d). After agroinfiltration, genes from Cluster I were strongly induced in CoVLP samples at late time points (Figure 5c), concomitant with accumulation of the S protein (Hamel *et al*., 2025). In fact, *NbASN1* and *NbASN2* were among the most strongly induced genes in CoVLP samples at 6 DPI, in accordance with proteomics data at 6 DPI confirming an accumulation rate of NbASN1 in CoVLP samples greater than in P19 samples (Hamel *et al*., 2025). By contrast, the genes from Cluster II were quickly repressed in both P19 and CoVLP samples, an effect more pronounced with the S protein expressed in leaf tissues (Figure 5d). No expression was detected for the sixth *ASN* gene, *NbASN3* (Niben101Scf11860g00012) (see asterisk in Figure 5b), suggesting this gene not to be expressed at detectable levels in collected leaf samples used for transcriptomics analyses. In light of such contrasted expression profiles, we propose that the *ASN* genes of Cluster I are involved in senescence-related reallocation of N resources, and the *ASN* genes of Cluster II in the biosynthesis of asparagine during vegetative growth (Figure 5a).

**Figure 5.**
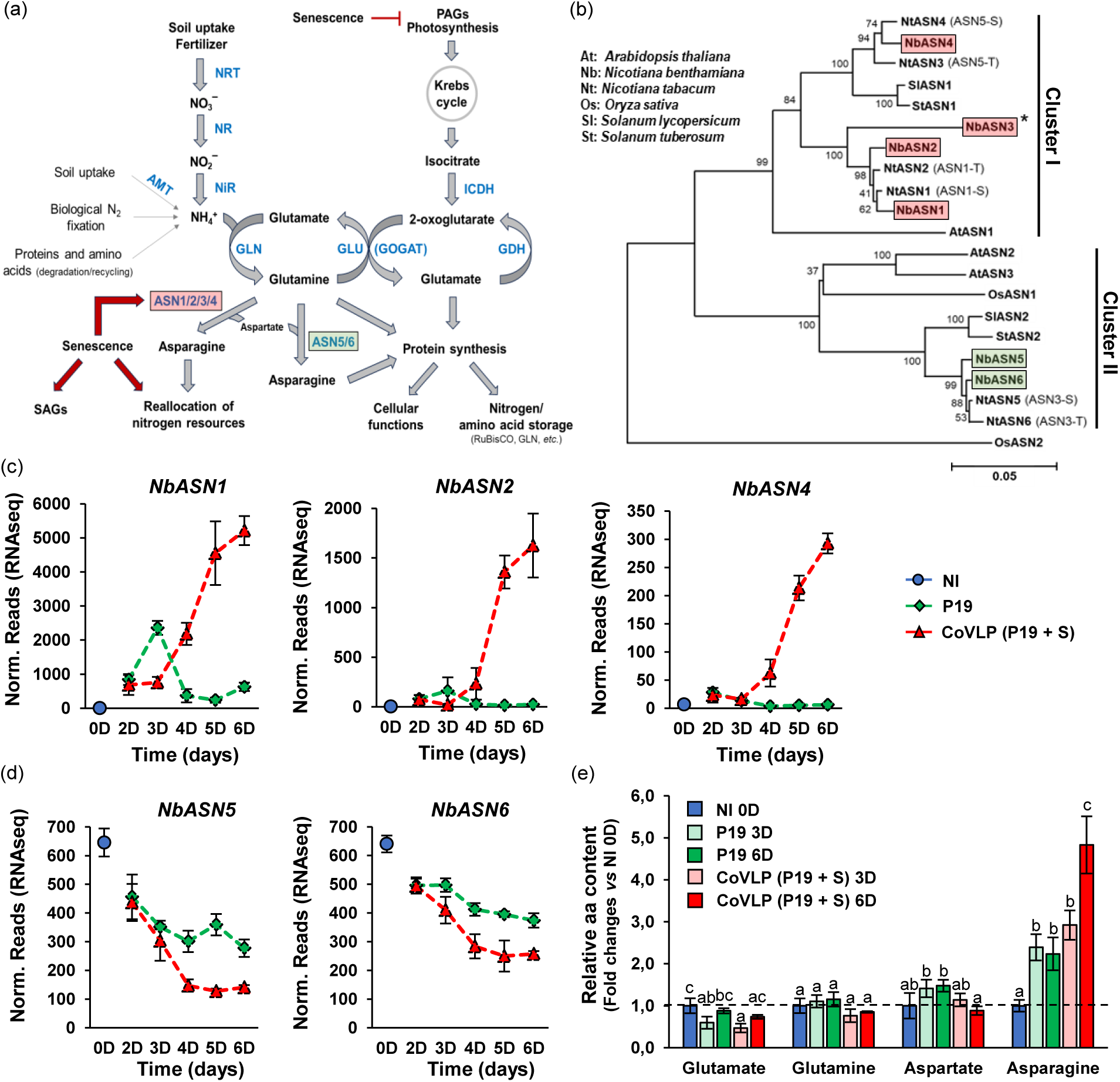
Phylogeny, regulation, and activity of NbASNs. (a) Assimilation of nitrogen (N) and effects of senescence on reallocation of N resources. Inorganic N is incorporated in amino acids as depicted. Metabolites are shown in black, proteins in blue. Adapted from Lu *et al*., 2016. Abbreviations: ammonium transporter (AMT), asparagine synthetase (ASN), glutamate dehydrogenase (GDH), glutamate synthase (GLU), glutamine 2-oxoglutarate aminotransferase (GOGAT), glutamine synthetase (GLN), isocitrate dehydrogenase (ICDH), nitrate reductase (NR), nitrate transporter (NRT), nitrite reductase (NiR). (b) Phylogeny of ASNs from various plant species. The *N. benthamiana* genome was searched using AtASNs as queries. Full-length protein sequences retrieved were aligned with ClustalW, using OsASN2 from rice as an outgroup. Resulting alignments were submitted to the MEGA5 software and a neighbor-joining tree derived from 5,000 replicates was created. Bootstrap values are indicated on the node of each branch. Senescence-related NbASNs are boxed in red, constitutive NbASNs in green. The asterisk (*) denotes lack of corresponding *NbASN3* gene expression in leaves that were harvested. For tobacco ASNs (NtASNs), alternative nomenclature is shown in parenthesis (Bovet *et al*., 2019). Expression profiles of senescence-related (c) or constitutive (d) *NbASNs*. For each time point in days (D) post-infiltration, RNAseq results are expressed in normalized (norm.) numbers of reads ± sd. (e) Relative amino acid (aa) contents at various time points. Amino acid contents of NI samples at 0 DPI were arbitrarily set at one-fold (dashed line). Groups that do not share the same letter are statistically different. Conditions are as follows: NI: non-infiltrated samples (blue); P19: agroinfiltrated samples expressing P19 only (green); CoVLP: agroinfiltrated samples co-expressing P19 and recombinant S protein (red).

Curing is used to induce senescence in tobacco leaves, leading to altered concentrations of the four amino acids glutamate, glutamine, aspartate, and asparagine, the latter being the most abundant (Bovet *et al*., 2019). To confirm a functional link between the upregulation of ASN-encoding genes and an eventual increase of the asparagine content in CoVLP-producing leaves, we determined the relative concentrations of glutamate, glutamine, aspartate and asparagine in leaf samples. The four amino acids were assayed by mass spectrometry (MS) in P19 and CoVLP samples collected at 3 and 6 DPI, arbitrarily setting their relative contents in NI samples at 0 DPI to one-fold (dashed line in Figure 5e). For glutamate, glutamine, and aspartate, no change of high magnitude was observed in P19 and CoVLP samples at 3 or 6 DPI. By contrast, asparagine content at 3 DPI was well over two-fold the asparagine content at 0 DPI for both P19 and CoVLP samples. Unlike in P19 samples, asparagine content further increased in CoVLP samples between 3 and 6 DPI, reaching an approximate five-fold increase compared to the baseline at 0 DPI (Figure 5e). This increase in asparagine concentration was consistent with accelerating accumulation of the S protein during this period (Hamel *et al*., 2025). Altogether, these observations formally confirmed a link between asparagine accumulation and *ASN* gene upregulation in the CoVLP samples, again strengthening the idea of a senescence-related, N remobilization-promoting effect of the CoVLPs in *N. benthamiana* leaves.

### Downregulation of genes involved in N assimilation

The regulation of *ASN* genes and increased asparagine content in CoVLP samples prompted us to look for an eventual downregulation of NH_4_^+^ assimilation through the GLN/GLU (GOGAT) cycle (Table S1). This pathway is key to the integration of inorganic N in the primary metabolism, notably as the main source of free amino acids for protein biosynthesis (Figure 5a). In the conditions examined, several *GLN* genes were differentially expressed. *NbGLN2a* (Niben101Scf02330g00012) and *NbGLN2b* (Niben101Scf00952g03003), two close homologs of Arabidopsis chloroplastic GLN-encoding gene *AtGLN2* (At5g35630), were gradually downregulated in P19 samples, and even more so in CoVLP samples (Figure 6a). In C3 plants such as *N. benthamiana*, chloroplastic GLNs are the predominant isoforms of GLN (Masclaux-Daubresse *et al*., 2006). By contrast, *NbGLN1b* (Niben101Scf01765g02012) was strongly induced, and its close homolog *NbGLN1a* (Niben101Scf00345g01021) slightly induced, at late time points in CoVLP samples (Figure 6b). *AtGLN1.3* (At3g17820), highly similar to *NbGLN1a*, and *AtGLN1.1* (At5g37600), highly similar to *NbGLN1b*, encode cytosolic GLNs induced during leaf senescence in Arabidopsis (Breeze *et al*., 2011). Upregulation of cytosolic GLNs is thought to facilitate N reassimilation following depletion of the chloroplastic GLN2 (Masclaux-Daubresse *et al*., 2010; Havé *et al*., 2017). RNAseq finally showed the gene *NbGLN1c* (Niben101Scf00824g00002) to be similarly repressed in both P19 and CoVLP samples (Figure 6b). Also encoding a cytosolic GLN, this gene was however expressed at much lower levels than the other *GLN* genes here identified (Figure 6a,b).

**Figure 6.**
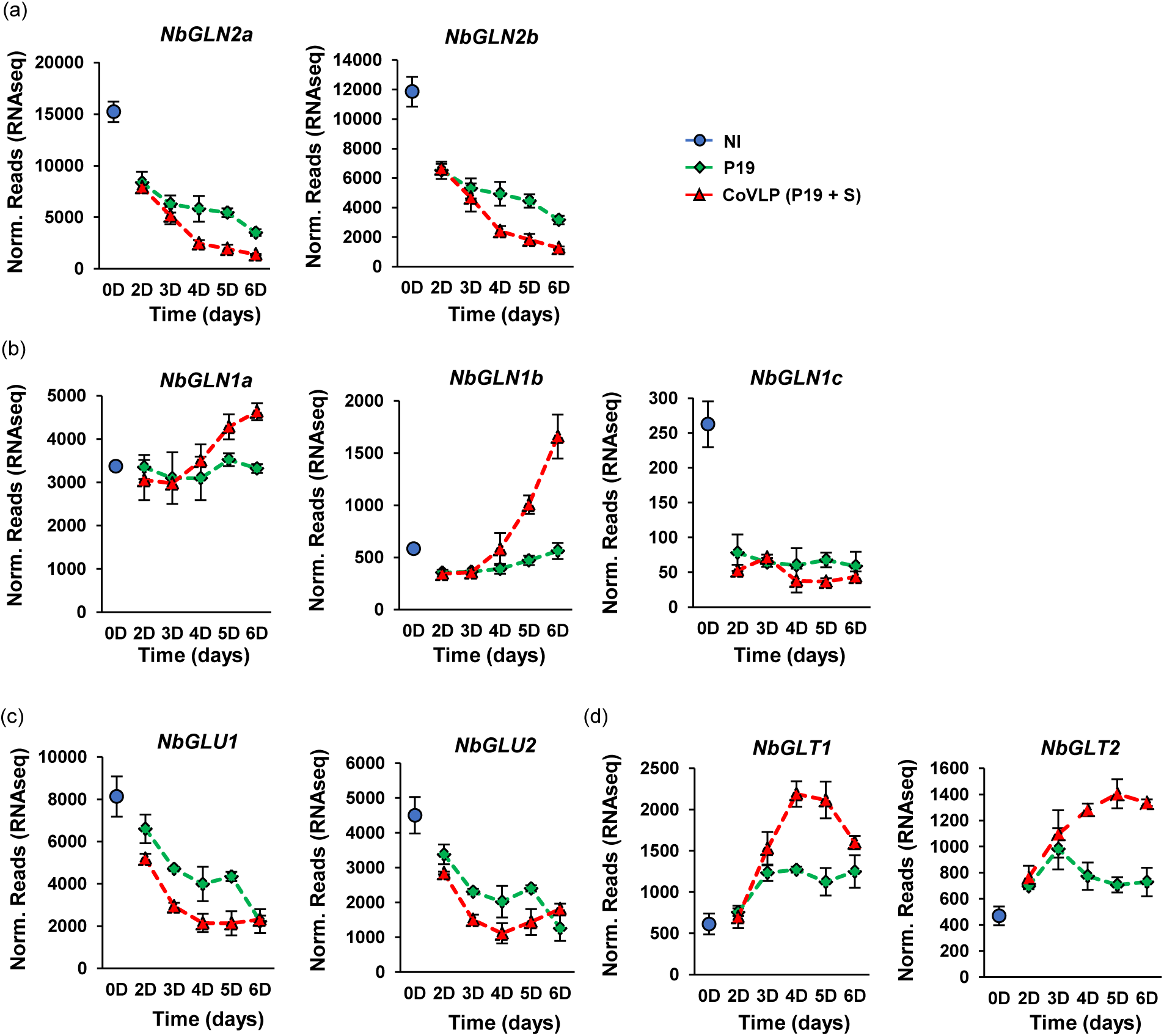
Expression of genes involved in N assimilation. Expression profiles of genes encoding chloroplastic GLNs (a), cytosolic GLNs (b), GLUs/Fd-GOGATs (c), and GLTs/NADH-GOGATs (d). For each time point in days (D) post-infiltration, RNAseq results are expressed in normalized (norm.) numbers of reads ± sd. Conditions are as follows: NI: non-infiltrated samples (blue); P19: agroinfiltrated samples expressing P19 only (green); CoVLP: agroinfiltrated samples co-expressing P19 and recombinant S protein (red).

Similar to *GLN g*enes, the most highly expressed *GLU/GOGAT* genes in leaves were downregulated after agroinfiltration (Table S1; Figure 6c,d). More specifically, chloroplastic ferredoxin-dependent glutamate synthase (GLU/Fd-GOGAT) gene *NbGLU1* (Niben101Scf04198g01001), similar to Arabidopsis *AtGLU1* (At5g04140), and its close homolog *NbGLU2* (Niben101Scf12966g00016), similar to *AtGLU2* (At2g41220), were gradually repressed in both P19 and CoVLP samples, with again more pronounced effects in the latter case (Figure 6c). By contrast, transcripts for nicotinamide adenine dinucleotide-dependent glutamate synthase (GLT/NADH-GOGAT) genes *NbGLT1* (Niben101Scf18214g00004) and *NbGLT2* (Niben101Scf02290g02021), two close homologs of Arabidopsis *AtGLT1* (At5g53460), were gradually induced in CoVLP samples, and to a lower extent in P19 samples, until reaching a maximum at 4 or 5 DPI (Figure 6d).

### Downregulation of genes involved in N reduction

Ammonium, used as a substrate by the GLN/GLU (GOGAT) cycle, is absorbed directly from the soil solution, or produced following NO_3_^−^ absorption and reduction through the sequential action of NR and NIR (Figure 5a). To investigate the impact of foreign protein accumulation on N reduction, we profiled the expression of genes encoding these N-reducing enzymes. Search of the *N. benthamiana* genome revealed the occurrence of two *NiR* genes, *NbNiR1* (Niben101Scf07103g03016) and *NbNiR2* (Niben101Scf02408g02009). RNAseq revealed both *NiR* genes to be downregulated early after agroinfiltration in both P19 and CoVLP samples (Figure 7a). As expression time progressed, the repression of *NbNiR1* and *NbNiR2* became more pronounced when CoVLPs were produced.

**Figure 7.**
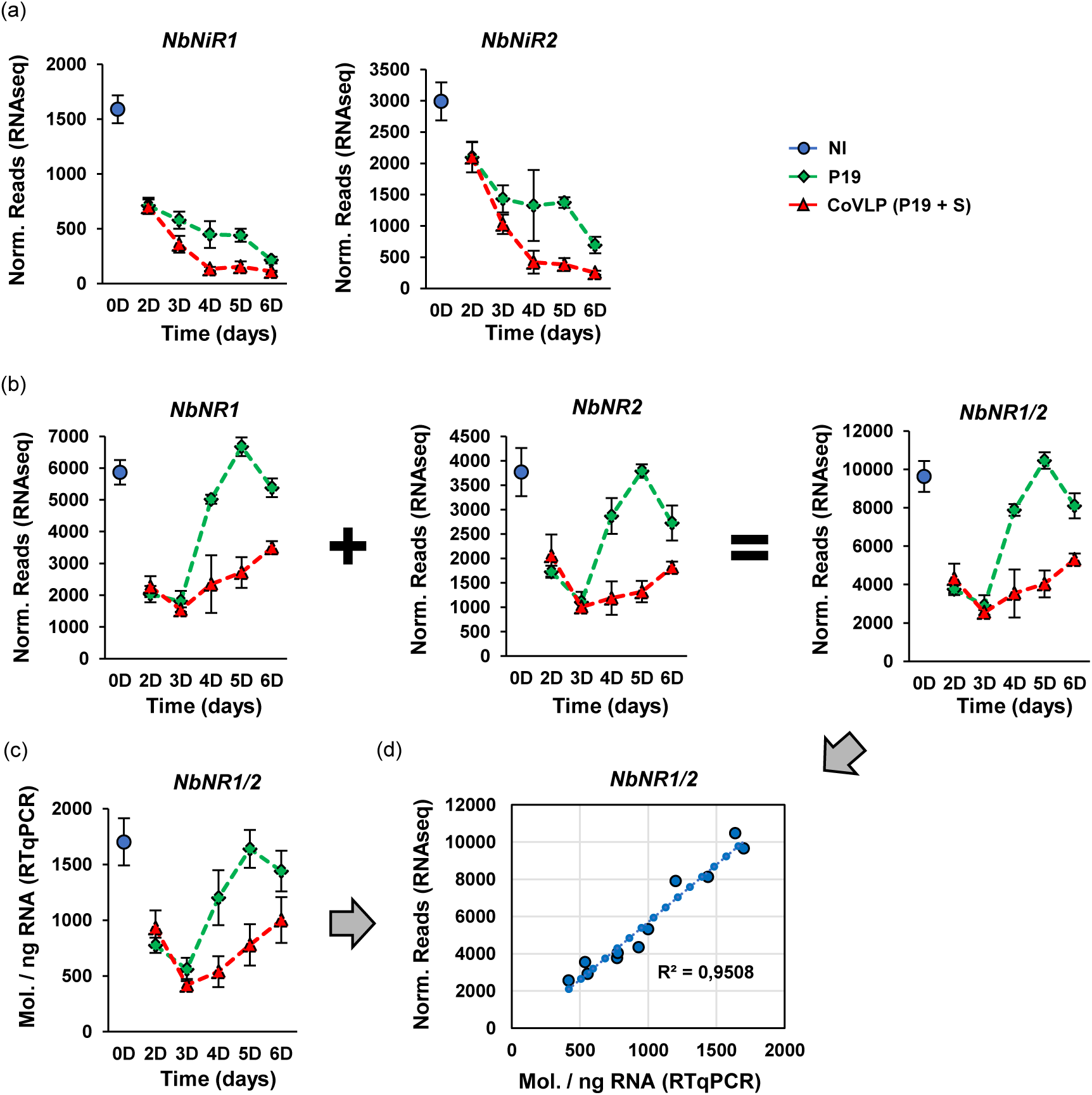
Expression of genes involved in reduction of NO_3_^−^ to NH_4_^+^. (a) Expression profiles of the two *NiR* genes of *N. benthamiana*. (b) Also shown are the expression profiles of *NR* genes *NbNR1* and *NbNR2*, or the combined expression of both genes (*NbNR1/2*) after the sum (+) of RNAseq reads. For each time point in days (D) post-infiltration, RNAseq results are expressed in normalized (norm.) numbers of reads ± sd. (c) Using primers specific to *NbNR1* and *NbNR2* transcripts, combined *NR* gene expression was also assessed by RTqPCR. Results are expressed in numbers of transcript molecules (mol.) per ng of RNA ± se. (d) Using RNAseq and RTqPCR results, linear regression and deduced coefficient of determination (R^2^) were deduced to highlight correlation between data from the two independent methods. Conditions are as follows: NI: non-infiltrated samples (blue); P19: agroinfiltrated samples expressing P19 only (green); CoVLP: agroinfiltrated samples co-expressing P19 and recombinant S protein (red).

Search of the *N. benthamiana* genome also revealed the occurrence of two *NR* genes, hereafter referred to as *NbNR1* (Niben101Scf01735g00005) and *NbNR2* (Niben101Scf34590g00006). Overall, these two genes exhibited very similar expression profiles, characterized by a rapid transcript depletion in both P19 and CoVLP samples at 2 and 3 DPI. Following this early repression, *NR* gene expression fully recovered at later time points in P19 samples, a recovery matched only in part in CoVLP samples (Figure 7b). RTqPCR assays with primers specific to *NbNR1* and *NbNR2* transcripts closely matched these observations (Figure 7c,d), again confirming the validity of our RNAseq datasets (Hamel *et al*., 2025).

### Deregulated NR as a helper protein to increase S protein accumulation in leaves

Given the SAG-inducing, CP-promoting, ASN-activating, and N assimilation-repressing effects of CoVLP accumulation, we concluded that the physiological context established in leaf cells upon S protein expression was likely to restrain the production of recombinant proteins. Based on this, we hypothesized that developing a strategy to sustain N reduction in leaf cells after agroinfiltration would help mitigate the negative impact of these molecular responses on protein biosynthesis, allowing for an increased S protein yield in leaf tissues. To test this, the *NbNR1* gene was cloned so it could be co-expressed as a helper protein along with the CoVLP construct. To maximize NR efficiency in agroinfiltrated plants over a 24-h cycle, a constitutively active, light-insensitive version of *NbNR1* was produced to prevent phosphorylation-mediated inactivation of NbNR1 in the dark (Douglas *et al*., 1995; Bachmann *et al*., 1996; Su *et al*., 1996). In Arabidopsis, the phosphorylation site for dark-induced inactivation corresponds to residue Ser-537 in AtNR1 (At1g77760) and to residue Ser-534 in AtNR2 (At1g37130) (Figure 8a). As this regulatory site corresponds to Ser-523 in NRs of tobacco and *N. benthamiana* (Figure 8a) (Lu *et al*., 2021), and considering the confirmed role of this residue for light responsiveness in tobacco NRs (Lillo *et al*., 2003), a variant of NbNR1 was created by substituting the Ser-523 residue for an aspartate, to generate a light-insensitive, deregulated form of the enzyme thereafter referred to as NbNR1^S523D^ (Figure 8a).

**Figure 8.**
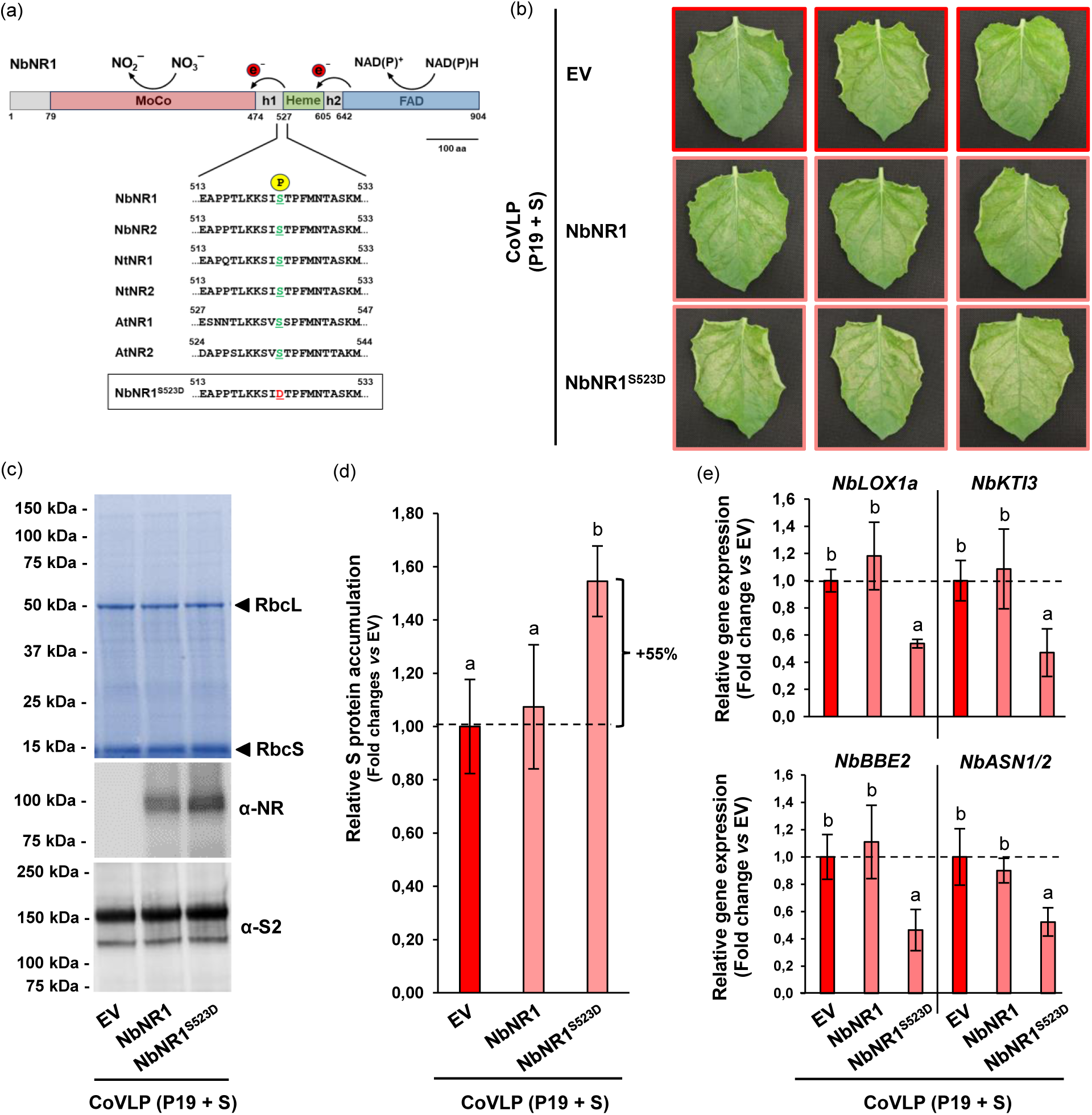
Co-expression of NbNR1^S523D^ increases S protein accumulation. (a) Topology of NbNR1 and model of its enzymatic activity. NRs have three functional domains, namely a molybdenum cofactor (MoCo) domain (red), an iron-binding heme domain (green), and a flavin adenine dinucleotide (FAD) domain (blue). Hinge (h) regions separate functional domains. To produce nitrite (NO_2_^-^), redox centers transfer two electrons (e-) from NADPH to nitrate (NO_3_^-^). Also conserved is a serine residue phosphorylated (P) to control dark inactivation of NR. A serine to aspartate substitution abolishes dark inactivation and results in constitutive activation of NR. Effects of helper proteins was evaluated by co-expressing P19 and the S protein with either NbNR1, NbNR1^S523D^, or an empty vector (EV) control. (b) Stress symptoms observed on representative leaves of each condition harvested at 6 DPI. Based on primary (P) leaf numbering (Hamel *et al*., 2025), pictures either highlight P9 or P10 leaves. (c) Total protein extracts following SDS-PAGE and Coomassie blue-staining (upper panel). Arrows highlight RuBisCO small (RbcS) and large (RbcL) subunits). Western blots depict accumulation of NRs (middle panel), or of the recombinant S protein (lower panel). (d) Relative S protein accumulation as measured by standardized capillary western immunoassays. S protein accumulation of CoVLP + EV samples was arbitrarily set at one-fold (dashed line). (e) Relative expression of CoVLP-induced genes as measured by RTqPCR. For each gene, transcript level of CoVLP + EV samples was arbitrarily set at one-fold. Error bars reflect the se. Groups that do not share the same letter are statistically different.

Co-expression experiments were conducted to assess the impact of NbNR1^S523D^ on S protein expression in agroinfiltrated leaves, using the wild-type enzyme for comparison purposes and an empty vector (EV) as a negative control. At 6 DPI, the intensity of leaf necrotic symptoms, characteristic of S protein expression (Hamel *et al*., 2025), was comparable in P9 and P10 leaves co-expressing CoVLPs along with wild-type NbNR1 or the EV control (Figure 8b). By comparison, the symptoms were more intense in leaves co-expressing the CoVLPs along with NbNR1^S523D^, as observed by more advanced leaf necrosis and curling of leaf blade edges (Figure 8b). Total protein extracts were also prepared using leaf biomass from whole plant shoots to estimate relative S protein accumulation by SDS-PAGE, western blotting, and standardized capillary western immunoassays. Coomassie blue staining indicated no obvious change in the overall patterns of protein extracts prepared from the three tested conditions (upper panel of Figure 8c). This analysis also confirmed that protein integrity had been maintained in the leaf extracts, including those from plants harbouring the most symptomatic leaves (Figure 8b). A western blot analysis with primary antibodies specific to conserved regions of plant NRs was then performed to confirm the presence of NbNR1 or NbNR1^S523D^ in the corresponding extracts (middle panel of Figure 8c). No signal was detected in CoVLP samples lacking a helper protein, suggesting the endogenous level of NR protein to be very low after 6 days of CoVLP production, consistent with the depletion of *NbNR1* and *NbNR2* transcripts in CoVLP samples at 6 DPI (Figure 7b,c). By comparison, clear signals were detected in CoVLP samples co-expressing NbNR1 or NbNR1^S523D^, with a more intense signal detected for the mutated enzyme. Together, these results indicated that both NbNR1 and NbNR1^S523D^ were overexpressed following agroinfiltration, and that substituting residue Ser-523 for an aspartate resulted in higher accumulation and/or stability of the mutated enzyme in leaf tissues compared to the original light-sensitive version.

To monitor S protein accumulation in the same protein samples as above, western blots were also produced using primary antibodies specific to the S2 domain of the SARS-CoV-2 S protein (lower panel of Figure 8c). Intense signals were detected in all tested conditions, indicating that co-expression of the helper proteins did not impair S protein accumulation in the plants. Standardized capillary western immunoassays were also performed to evaluate relative S protein levels with more accuracy, assigning an arbitrary value of 1.00 to the samples prepared from control plants lacking helper protein co-expression (dashed line in Figure 8d). Despite sustained expression, NbNR1 had no beneficial effect on S protein content in leaf extracts compared to the EV control. On the opposite, S protein accumulation was increased by ∼55% in CoVLP samples co-expressing NbNR1^S523D^ compared to the baseline where no helper protein was expressed (Figure 8d). This observation was consistent with intensity of the leaf symptoms, which were more advanced when NbNR1^S523D^ was co-expressed (Figure 8b).

To further assess the impact of NbNR1^S523D^ co-expression *in planta*, RTqPCR measurements were next performed with primers specific to several defense gene markers. For those analyses, RNA extracts employed were derived from the pooled leaf biomass of whole plant shoots, which contained a mixture of symptomatic and asymptomatic leaves used to quantify the S protein at the whole plant scale (Figure 8c,d). Examined defense gene markers included *NbLOX1a* (Niben101Scf01434g03006), *NbKTI3* (Niben101Scf06424g00003), and *NbBBE2* (Niben101Scf00944g01001), reported to be strongly and specifically induced at late time points in CoVLP samples (Hamel *et al*., 2025). Also analysed were the senescence-inducible genes *NbASN1* and *NbASN2*, upregulated at late time points in CoVLP samples (Figure 5c). Relative expression levels were determined for each gene marker by arbitrarily setting at 1.00 the expression level detected in the CoVLP + EV control samples (dashed lines in Figure 8e). For all tested markers, expression levels in CoVLP samples co-expressing wild-type NbNR1 were similar to expression levels in CoVLP + EV control samples, in contrast with the reduced transcript numbers in CoVLP samples co-expressing NbNR1^S523D^. Very similar trends were observed for CoVLP-inducible genes *NbPDF1* (Niben101Scf17290g01005), *NbPPO1* (Niben101Scf00180g08002), and *NbPPO3* (Niben101Scf04384g02014) (Figure S2a) (Hamel *et al*., 2025). By comparison, no changes in transcript levels were noted for other defense markers such as the pathogen-inducible genes *NbPR1a* (Niben101Scf01999g07002) and *NbPR1b* (Niben101Scf00107g03008), or the sphingolipid biosynthesis genes *NbIPCS1* (Niben101Scf01406g01001) and *NbIPCS2* (Niben101Scf04368g00005) (Figure S2b) (Hamel *et al*., 2025). Overall, these observations pointed to a positive, promoting effect of NbNR1^S523D^ on CoVLP accumulation in leaf tissue, likely explained by an improved potential for NO_3_^−^ reduction and a concomitant downregulating effect on some CoVLP-inducible defense-and senescence-related genes.

## Discussion

Our aim in this study was to document the possible inducing effect of S protein expression and CoVLP accumulation on senescence-related processes in agroinfiltrated leaves of *N. benthamiana* plants used as biofactories for COVID-19 vaccine production. In a recent study on the molecular responses of *N. benthamiana* leaf cells expressing H5, an influenza virus hemagglutinin that also induces the formation of enveloped VLPs, we highlighted the upregulation of gene markers for senescence in leaf tissues, including *NbORE1*, *NbSGR1*, Cluster I *NbASNs*, and key genes of the ethylene biosynthetic pathway (Hamel *et al*., 2024). In support to these initial observations, we here provide several lines of evidence confirming the induction of senescence upon CoVLP accumulation. Senescence marks observed comprised the upregulation of several senescence-related genes, the downregulation of primary metabolism-associated genes, and the production of asparagine for N recycling in alternative organs and tissues. From these varied findings, we conclude that leaf senescence represents, along with several immune responses (Hamel *et al*., 2024, 2025), a central feature of the host plant’s response to the production of enveloped VLPs following agroinfiltration and expression of virus surface proteins such as the influenza H5 and the SARS-CoV-2 S proteins. Considering that H5 and the S protein share very limited homology, our findings also suggest that the activation of senescence-related processes in VLP-producing leaves is triggered by the accumulation of enveloped nanoparticles rather than by the identity of the viral proteins at the surface of these nanoparticles.

Except for CP activity and accumulation of the SAG12-like proteases, the senescence-related responses here described were observed in young, actively growing leaves located in the upper part of the plants. These leaves were selected for biomass harvesting given their high efficiency at expressing the recombinant S protein (Hamel *et al*., 2025) and their relevance as a model to characterize growth-immunity trade-offs in the host plant upon agroinfiltration (Hamel *et al*., 2024). Detecting the characteristic molecular signatures of senescence at such an early stage of leaf development may seem surprising at first sight, but not so much considering the accelerating effect of environmental stresses of this process (Schippers *et al*., 2015; Woo *et al*., 2019) and the artificial, inherently stressful nature of recombinant protein production using the leaf agroinfiltration system. Possible sources of stress under our conditions included mechanical stress caused by the infiltration procedure itself, bacterium perception by leaf cells, and temporal accumulation of the P19 and S proteins, the latter enabling formation of nanoparticles budding from the PM of plant cells. We suggest, at this point, this intricate combination of stress cues to have triggered senescence-related processes despite the active metabolic status and young physiological age of the leaves selectively collected for analysis (Hamel *et al*., 2025).

Molecular responses in leaf tissues expressing the S protein included the upregulation of *NAC* genes that encode homologs of senescence-promoting NACs in other plant species (Podzimska-Sroka *et al*., 2015; Li *et al*., 2018; Nagahage *et al*., 2018; Chen *et al*., 2023). *WRKY* genes were also upregulated (Hamel *et al*., 2025), many of them showing similarity to Arabidopsis *WRKYs* that promote leaf senescence (Robatzek and Somssich, 2001; Guo *et al*., 2004; Zhou *et al*., 2011; Niu *et al*., 2020). Upregulated *NbNACs* and *NbWRKYs* encompassed early and late genes, categorized based on the timing of their expression pattern compared to the accumulation of the S protein, as also proposed for a number of defense-related TF genes (Hamel *et al*., 2025). The onset of two induction waves for TF-encoding genes suggests a certain hierarchy among senescence-promoting TFs, presumably to fine-tune the activation of early and late mechanisms orchestrating different phases of the leaf senescence process. Complex relationships among transcriptional regulators during leaf senescence have been documented in Arabidopsis, such as for AtNAC2 and AtNAC81, both interacting with the promoter of *AtORE1* to favour its expression (Garapati *et al*., 2015; Nagahage *et al*., 2018). During senescence, *AtORE1* is also regulated by a signalling cascade comprising *ETHYLENE-INSENSITIVE 2* (*AtEIN2*; At5g03280), *ETHYLENE-INSENSITIVE 3* (*AtEIN3*; At3g20770), and *microRNA164* (*miR164*) (Kim *et al*., 2009; Li *et al*., 2013). While TF AtEIN3 represses the accumulation of *miR164*, which itself works as a repressor of *AtORE1* expression (Li *et al*., 2013), it also binds the promoters of *AtORE1* and *AtNAC29* to induce synthesis of the corresponding TFs (Kim *et al*., 2014).

As close homologs of all these senescence-promoting *NACs* were induced in leaves upon S protein expression, we took a closer look at the expression patterns of potential upstream components eventually playing a role in this important signalling pathway. The two closest homologs of *AtEIN2*, here termed *NbEIN2a* (Niben101Scf23355g00004) and *NbEIN2b* (Niben101Scf04548g00001), showed unchanged transcript numbers during S protein expression (Figure S3a) or no transcript at all in leaves that were sampled, respectively. By contrast, a *N. benthamiana* homolog of *AtEIN3*, *NbEIN3a* (Niben101Scf00304g01003), was severely repressed throughout the expression phase in both P19 and CoVLP samples, unlike other close homologs *NbEIN3b* (Niben101Scf02891g00008) and *NbEIN3c* (Niben101Scf17776g00004) being induced already at 2 DPI and even more at later time points in CoVLP samples (Table S1; Figure S3b). While suggesting the involvement of additional regulatory genes for the activation of leaf senescence in *N. benthamiana*, these observations also underline the striking complexity of senescence-related signaling in leaves following agroinfiltration and the likely role of multigene families in the regulation of this process. Senescence-related genes induced following agroinfiltration also included SAGs with varied functions, as well as other sets of genes involved in chlorophyll degradation, senescence-related PCD, ethylene biosynthesis, and senescence-related proteolysis. Further studies will be required to identify transcriptional regulators controlling the expression of these senescence markers but current knowledge in Arabidopsis suggests a role for at least some components of the same TF regulatory networks. For instance, AtNAC87, AtORE1, and AtEIN3 induce the expression of genes involved in chlorophyll degradation (Qiu *et al*., 2015; Chen *et al*., 2023), including *AtSGR1* and *AtPaO*, two close homologs of *NbSGR1* and *NbPaO1* induced in leaves upon S protein expression. Likewise, AtNAC81 and AtORE1 promote the senescence-related accumulation of ethylene through enhanced expression of ethylene biosynthesis genes (Qiu *et al*., 2015; Peng *et al*., 2022), including *AtACS2* (At1g01480), a close homolog of *NbACS3* again strongly induced in CoVLP samples. These inferences, although requiring functional validation in *N. benthamiana*, suggest that at least some of the TF-encoding genes upregulated during CoVLP production belong to a common leaf senescence-promoting transcriptional network in plants. From a molecular farming standpoint, deciphering transcriptional regulatory networks that control defense-and senescence-related genes is of great significance, given their potential to interfere with primary metabolic functions in plant cell biofactories, and in turn to affect host plant fitness and efficiency to produce proteins.

As evidenced by the downregulation of several PAGs, senescence-related responses following agroinfiltration and CoVLP production appear to interfere with important chloroplastic functions, including light capture and synthesis of photosynthetic pigments. In several cases, the expression of PAGs is promoted by TFs of the GLK family (Waters *et al*., 2009), shown here to be repressed in both P19 and CoVLP samples. As master transcriptional regulators, GLKs are controlled at several levels in leaf cells. In Arabidopsis, AtORE1 for instance antagonizes the transcriptional activity of GLKs through direct protein-protein interactions (Rauf *et al*., 2013). At the transcriptional level, AtNAC2 shifts the physiological balance of plant cells towards leaf senescence by activating the expression of *AtORE1* while at the same time inhibiting the expression of *AtGLK1* (Garapati *et al*., 2015). On the opposite, MYB-related TF CIRCADIAN CLOCK-ASSOCIATED 1 (AtCCA1; At2g46830) prevents the activation of leaf senescence by activating the expression of *AtGLK2* while inhibiting the expression of *AtORE1* (Song *et al*., 2018). In *N. benthamiana*, the closest homolog of AtCCA1, *NbCCA1* (Niben101Scf02026g01002), was severely repressed in both P19 and CoVLP samples (Table S1; Figure S3c), again consistent with the idea that foreign protein expression activates senescence in agroinfiltrated leaves. Based on this, we conclude that by repressing chloroplast-associated genes, recombinant protein expression generates a cellular environment to some extent unfavorable to the maintenance of chloroplastic functions, another hallmark of leaf senescence.

In addition to PAGs, transcriptomics showed CoVLP accumulation to inhibit the expression of several genes involved in N reduction and N assimilation. These two metabolic functions are essential in plants to sustain sufficient amino acid pools for protein biosynthesis, and the downregulation of either functions in agroinfiltrated leaves likely represents a constraint to their productivity as protein biofactories. Most key genes driving N reduction to NH_4_^+^, or NH_4_^+^ assimilation in amino acids, were strongly repressed during CoVLP production, including *NRs*, *NiRs*, and chloroplastic *GLNs*. This was in sharp contrast with senescence-related *ASN* genes being strongly upregulated, and with the consistent accumulation of asparagine in CoVLP samples at 6 DPI. Together, these findings further point to the onset of senescence-related mechanisms upon CoVLP accumulation, which not only causes a slowdown of primary metabolic functions, but also premature recycling of N resources from tissues that should otherwise not senesce considering their young stage of development. In practice, these observations suggest that cellular responses to recombinant protein accumulation impose ceiling in accumulation of CoVLPs by limiting N availability. Despite production rates sufficient to sustain the large-scale production of COVID-19 vaccines in *N. benthamiana* (Hager *et al*., 2022), deficient NO_3_^−^ reduction, and as a consequence a suboptimal source of NH_4_^+^ cations for N assimilation, thus represented an interesting target for improvement of S protein yields in leaf tissues.

Nitrate reduction has been the focus of several attempts to improve N use efficiency in plants, including via the constitutive overexpression of *NR* genes (Pathak *et al*., 2009; Masclaux-Daubresse *et al*., 2010). Although providing notable advantages under specific physiological conditions, the actual agronomical benefits of this approach remain unclear given the complex post-translational regulation of NR, including a rapid inactivation by phosphorylation in dark conditions. Accordingly, co-expression of *NbNR1* in agroinfiltrated leaves along with the *S* gene had little impact on S protein accumulation, while the co-expression of its constitutively active variant NbNR1^S523D^ increased S protein level by more than 50%. A first, likely explanation for such an improved yield would be a better availability of amino acids and the maintenance of a more favorable cellular environment for protein biosynthesis, despite immunity activation and the onset of senescence-related processes. At the whole plant scale, a second, non-exclusive explanation would be the indirect downregulating impact of NbNR1^S523D^ on defense-and senescence-related genes through a partial rebalancing of host cell primary metabolic functions towards anabolism. In a production context, further studies will be required to confirm the yield-promoting effect of NbNR1^S523D^ at the plant canopy level, as this effect might vary according to the age or physiological status of the leaves. Further studies will also be warranted to better elucidate the trade-offs established between primary metabolism, immunity responses, and leaf senescence in different parts of the canopy. Despite several questions remaining, data provided in this study significantly expand our understanding of plant responses to foreign protein expression, in addition to providing a genetic approach to increase S protein yields for COVID-19 vaccine development. It will now be interesting to see whether this approach also benefits the expression of other recombinant proteins commonly produced in plants, including influenza hemagglutinins and therapeutic antibodies.

## Materials and methods

### Seed germination and plant growth

Wild-type *N. benthamiana* plants were used for all experiments. Detailed procedures and conditions used for seed germination and plant growth are described in our accompanying manuscript (Hamel *et al*., 2025).

### Binary vector constructs

To express CoVLPs, a recombinant version of the SARS-CoV-2 S protein (original strain Wuhan-Hu-1) was employed. Details on source of the used sequence, modifications that were introduced and cloning in the final binary vector are provided in Hamel *et al*. (2025). To prevent silencing of the *S* transgene in leaves after agroinfiltration, the T-DNA region of the CoVLP construct also comprised the tomato bushy stunt virus (TBSV) suppressor of RNA silencing gene *P19*, under the control of *plastocyanin* promoter and terminator sequences. For P19 samples, a binary vector allowing expression of P19 only was used.

For genetic constructs allowing the co-expression of *NbNR1* and *NbNR1^S523D^*, full-length nucleotide sequence of *NbNR1* was retrieved from the genome assembly database of *N. benthamiana* (https://solgenomics.net/). The wild-type sequence was codon-optimized and the resulting template used to design a synthetic gene block (gBlock) (Integrated DNA Technologies). To produce *NbNR1^S523D^*, the same strategy as above was applied, except that a ‘TCA’ codon was replaced by a ‘GAT’ codon to encode an aspartate instead of a serine at position 523 of the corresponding NbNR1 protein. gBlocks were amplified by PCR and individually introduced using the In-Fusion cloning system (Clontech) in the T-DNA region of a customized pCAMBIA0380 binary vector previously linearized with restriction enzymes *SacII* and *StuI*. The expression of *NbNR1* and of *NbNR1^S523D^* was driven by a *2X35S* promoter from the cauliflower mosaic virus (CaMV). The expression cassette also comprised proprietary 5’-and 3’-untranslated regions (UTRs) developed to maximize mRNA stability and protein translation, as well as the *Agrobacterium nopaline synthase* (*NOS*) gene terminator. An empty binary vector was used as control for the co-expression experiments.

### Agrobacterium cultures and plant infiltration

All binary vectors were heat shock-transformed in *A. tumefaciens* strain AGL1. Details on handling of transformed bacteria, preparation of glycerol stocks, growth of bacterial cell cultures, and preparation of final inocula for agroinfiltration are provided in Hamel *et al*. (2025). For S protein expression, a final OD_600_ of 0.6 was used. For co-expression of S protein and helper proteins, separate bacterial cell cultures were prepared and mix prior to infiltration so final OD_600_ of 0.6 and 0.2 are reached, respectively. Vacuum infiltration was performed by placing whole plant shoots upside down in an airtight stainless-steel tank containing the appropriate bacterial suspension. To draw air out of the leaves, vacuum pressure was applied for 1 min, before pressure release to force the bacterial inoculum into the leaves.

### Transient protein expression and biomass harvesting

Recombinant protein accumulation was allowed to proceed in agroinfiltrated leaves for several days, as indicated. Protein expression took place in plant growth chambers using conditions described in Hamel *et al*. (2025). For sampling of biomass used for RNAseq, RTqPCR, proteomics, and quantification of amino acids, P leaves of similar developmental stage (P9 and P10) were selectively harvested using a leaf numbering approach described in Hamel *et al*. (2025). Freshly cut leaves without petiole were placed in pre-frozen 50 mL Falcon tubes, before flash freezing in liquid N. Each sample was made from six leaves collected on three randomly selected plants. For western blot on SAG12-like proteases, measurements of CP activity, and co-expression of helper proteins, primary and secondary leaves from whole plant shoots were manually detached, avoiding axillary stems and leaf petioles as much as possible. Biomass from three randomly selected plants was collected, placed in aluminum foil, and flash frozen in liquid N. For all samples, frozen biomass was stored at -80°C until ready for analysis. Foliar tissues were then ground and homogenized to powder in liquid N using pre-chilled mortars and pestles. For all experiments, the average results presented were obtained from at least three biological replicates.

### Protein extraction and quantification

Buffers and methods used for protein extraction and protein quantification in crude extracts are provided in Hamel *et al*. (2025).

### Western blotting and standardized capillary western immunoassays

Buffers and methods used for SDS-PAGE, protein transfer, and immunoblotting are provided in Hamel *et al*. (2025). Primary antibody dilutions were as follows: anti-SARS-CoV-2 Spike S2 polyclonal antibody (Novus Biologicals): 1/20,000; anti-SAG12 polyclonal antibody (Agrisera): 1/2,000; anti-NR polyclonal antibody (Agrisera): 1/2,000. Secondary antibody dilution was as follows: goat anti-rabbit (JIR): 1/10,000.

For relative quantification of the S protein, standardized capillary western immunoassays were performed using WES or JESS devices (ProteinSimple). Details on the preparation of standard curves, S protein controls, and samples to be analysed are provided in Hamel *et al*. (2025), along with the parameters employed to capture and analyse the resulting data. For S protein detection, an anti-SARS-CoV-2 Spike S2 polyclonal antibody (Novus Biologicals) and a goat anti-rabbit antibody (JIR) were used as primary and secondary antibodies, respectively.

### RNA extracts, RNAseq and RTqPCR analyses

Procedures used for extraction, quantification, qualification, and long-term storage of total RNA extracts are provided in Hamel *et al*. (2025). RNAseq analyses were performed to study global changes to the leaf transcriptomes, using RNA from three biological replicates of each experimental condition and time point. Methods employed for the preparation of cDNA libraries, Illumina sequencing, quality control assessments, and management of large-scale sequencing data are described in Hamel *et al*. (2025). For the profiling of individual genes, variance-stabilized read values were used. The complete set of RNAseq data used in this study has been deposited in the European Nucleotide Archive (ENA) database (ebi.ac.uk/ena/browser/home), under accession number PRJEB64341.

To validate RNAseq data, RTqPCR was performed on a variety of plant genes (Hamel *et al*., 2025), including *NbNR1* and *NbNR2* here presented as complete time series. Detailed methods and primer sequences for the RTqPCR analyses are available in Hamel *et al*. (2025).

### Large-scale proteomics

To study changes in protein abundance, total protein extracts from P19-and CoVLP-expressing leaves harvested at 6 DPI were used for iTRAQ labeling. Three biological replicates were used for each condition analysed. Methods used to extract and label proteins are described in Hamel *et al*. (2025).

### Measurement of the CP activity

The activity of CPs was monitored using substrate hydrolysis progress curves (Goulet *et al*., 2008) and the synthetic fluorogenic peptide *Z*-Phe-Arg-MCA (Pepnet), a preferred substrate for cathepsin L-like CPs. Leaf proteins were extracted using assay buffer (125 mM citrate, pH 6.2, 200 mM mannitol, 500 mM NaCl, and 25 mM EDTA), supplemented with 10 mM L-cysteine and 1% (w/v) PEG 8000. After clarification by centrifugation, protein extracts were diluted in assay buffer at ratios of 1/150, 1/300, and 1/600. To quantify CP activity, standard curves were made using recombinant human cathepsin L available commercially (Abcam). Based on the enzymatic unit (U) definition provided by the manufacturer, standard curves comprised seven points ranging from 1 mU to 300 mU. After dilution, a 100 µL volume of each standard curve point and of each sample dilution was loaded in white, opaque 96-well microplates (Pierce). To reach a final assay concentration of 8 µM, 100 µL of 16 µM *Z*-Phe-Arg-MCA was added to each well. Microplates were then gently shaken for 10 sec, before the first activity measurements were taken. Using the fluorescence mode of a SpectraMax M2 microplate reader (Molecular Devices), hydrolysis of the fluorogenic substrate was allowed to proceed for 10 min at room temperature, with readings performed every 25 sec. The excitation wavelength was set at 360 nm and the emission wavelength at 450 nm. Predetermined photo multiplicator setting of the device was set to medium. Once reaction time was completed, captured data was transferred to the Microsoft Excel software for final analysis.

### Amino acid quantification

For the quantification of amino acids, sample preparation and data acquisition were performed by metaSysX (https://www.metasysx.com). A protocol adapted from Salem *et al*. (2016) was followed to extract polar and semi-polar metabolites, lipids, proteins, and cell wall polymers from a single biomass. For each sample, 20 mg of ground leaf tissues was used. Metabolomic analyses were performed using a Waters ACQUITY Reversed Phase Ultra Performance Liquid Chromatography (RP-UPLC) device, coupled to a ThermoFisher Exactive mass spectrometer consisting of an electrospray ionization source (ESI) and an Orbitrap mass analyzer, as well as with an Agilent Technologies mass spectrometer consisting of an electron impact ionization source (EI) and a time of flight (TOF) mass analyzer. UPLC-MS measurements of the aqueous phase enabled for the detection of polar and semi-polar metabolites, including amino acids.

### Phylogenetic analyses

To better characterize CPs identified in this study, the genome of *N. benthamiana* was searched using full-length amino acid sequences of Arabidopsis AtCathB1, AtSAG2, AtSAG12, AtRD19a, or AtRD21a as queries. The predicted N-terminal SPs and pro-domains were manually removed from full-length protein sequences that were retrieved, before aligning the predicted mature protease sequences using ClustalW. To identify members of the ASN family, the genome of *N. benthamiana* was searched using the full-length amino acid sequences of Arabidopsis AtASN1 (At3g47340), AtASN2 (At5g65010), and AtASN3 (At5g10240) as queries. Full-length protein sequences retrieved were again aligned with ClustalW. For both CPs and ASNs, the alignment parameters were as follows: for pairwise alignments, 10.0 for gap opening and 0.1 for gap extension; for multiple alignments, 10.0 for gap opening and 0,2 for gap extension. The resulting alignments were submitted to the MEGA5 software and neighbor-joining trees derived from 5,000 replicates were generated. Bootstrap values are indicated on the node of each branch.

### Statistical analyses

For measurement of the CP activity and quantification of amino acids, statistical analyses were performed using Graph Pad Prism 10.1.0. Two-way ANOVA were used, with time points as row factors and conditions as column factors. Conditions were then analysed by post hoc Tukey’s multiple comparison tests with an alpha threshold of 0.05. For standardized capillary western immunoassays and RTqPCR analyses on samples expressing the helper proteins, statistical analyses were also performed using Graph Pad Prism 10.1.0. Groups were analysed using one-way ANOVA, followed by a post-hoc Tukey’s multiple comparison test with an alpha threshold of 0.05. Groups are labeled with a compact letter display. Groups that do not share the same letter are statistically different.

## Acknowledgments and funding

This work would not have been possible without the support of the Biomass Production and Research & Innovation teams of Medicago. Contributions included, but were not limited to, cloning of the gene constructs, inoculum preparation, plant cultivation, and leaf agroinfiltration. The authors also wish to acknowledge Cecilia Cheval, James Nicolas Duncan Battey, Lucien Bovet, and Simon Goepfert (current or former employees of Philip Morris International) for assistance with study design, sequencing data acquisition, and helpful discussions. Funding for this work was provided by Medicago Inc.

## Author contributions

L.P.H., M.A.D., and D.M. designed the research, supervised the project, and analysed the data. L.P.H. and M.A.C. drafted the manuscript and assembled the figures. P.O.L. managed production of the genetic constructs. L.P.H., F.P.G., R.T., M.E.P., and M.A.C. produced biomass and extracts used for transcriptomics and proteomics analyses. L.P.H., M.A.C., and F.P.G. managed RNAseq data and performed gene expression profiling. M.C.G., A.L., M.E.P., and M.A.C. developed the proteolytic activity assay and performed quantification of the CP activity. L.P.H. and M.C.G. performed phylogenetic analyses and protein sequence alignments. L.P.H. analysed data from the amino acid quantification. R.T. and M.E.P. performed RTqPCR analyses. M.E.P., F.P.G., R.T., M.A.C, and L.P.H. conducted the NR co-expression experiments. M.E.P. performed assays for recombinant S protein quantification and western blots on SAG12-like proteases and NRs. M.A.C., M.C.G., and F.P.G. performed statistical analyses. All authors read, helped to edit, and approved final version of the manuscript.

## Data availability statement

All data discussed in this study can be found in the manuscript and in the Supplementary Materials.

## Supporting information

Additional supporting information may be found online in the Supporting Information section at the end of the article.

**Figure S1.**
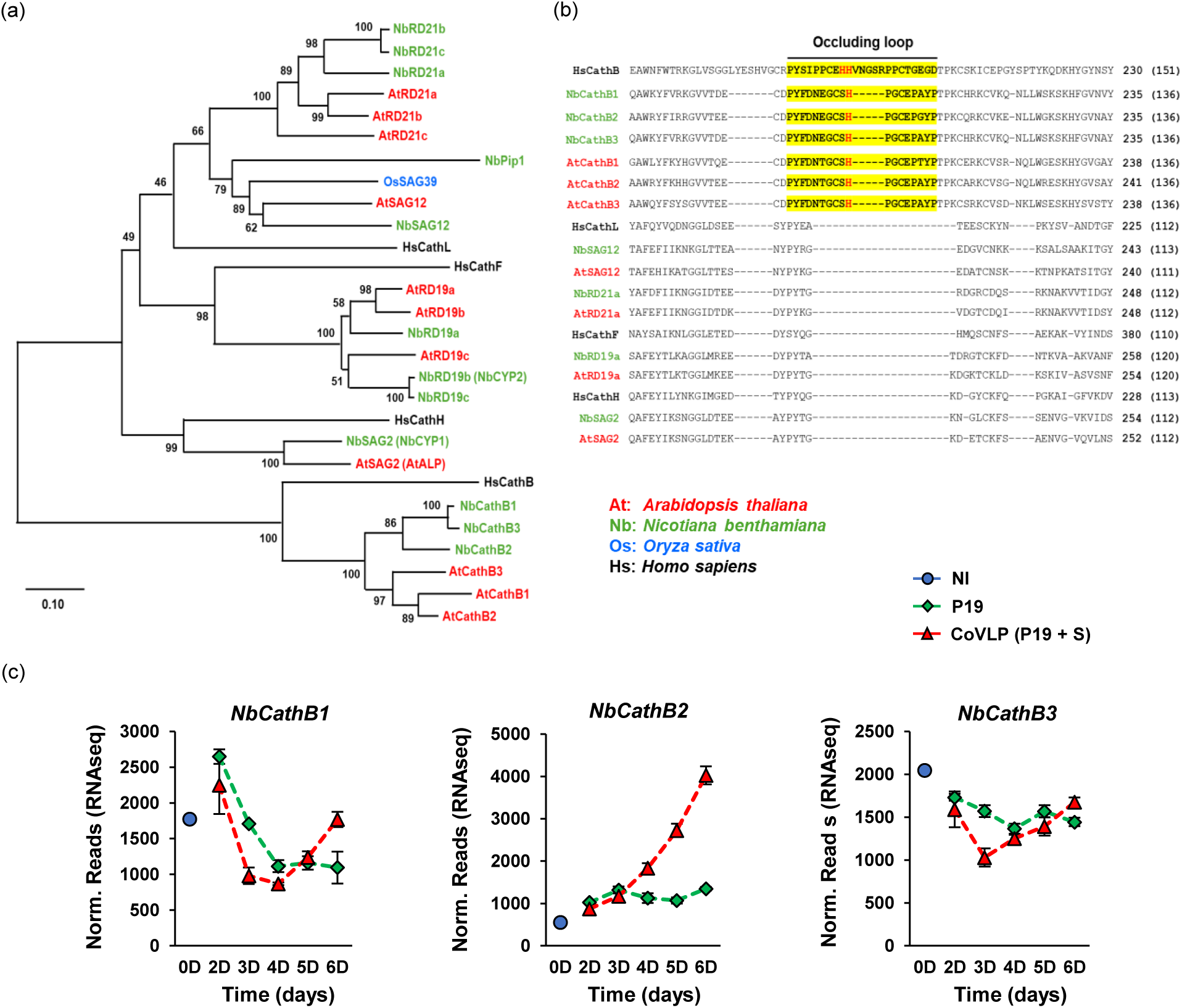
Classification of CPs and transcriptional profiles of *NbCathBs*. (a) Phylogeny of CPs from various plant species. The *N. benthamiana* genome was searched using amino acid sequences of AtCathB1, AtSAG2, AtSAG12, AtRD19a, or AtRD21a as queries. From full-length protein sequences that were retrieved, N-terminal SPs and pro-domains were removed so predicted mature proteases could be aligned with ClustalW. As additional group references, *Homo sapiens* cathepsin B (HsCathB; NP_001371643), cathepsin F (HsCathF; NP_003784), cathepsin H (HsCathH; NP_004381), and cathepsin L (HsCathL; NP_001244900) were included in the analysis. Resulting alignments were submitted to the MEGA5 software and a neighbor-joining tree derived from 5,000 replicates was created. Bootstrap values are indicated on the node of each branch. (b) Protein sequence alignment of various cathepsins from plants and human. The alignment section presented highlights a conserved CathB region that comprise an occluding loop absent in other cathepsin types (highlighted in bold yellow). Compared to HsCathB, occluding loops of plant CathBs are shorter and comprise only one histidine residue (highlighted in bold red). These histidines are strictly conserved as they are required for peptidase activity. Amino acids are depicted using their one-letter code. On the right, amino acid positions within full-length proteins are indicated and corresponding positions in mature proteases are in parenthesis. (c) Expression profiles of *NbCathB* genes identified by RNAseq. For each time point in days (D) post-infiltration, RNAseq results are expressed in normalized (norm.) numbers of reads ± sd. Conditions are as follows: NI: non-infiltrated samples (blue); P19: agroinfiltrated samples expressing P19 only (green); CoVLP: agroinfiltrated samples co-expressing P19 and recombinant S protein (red).

**Figure S2.**
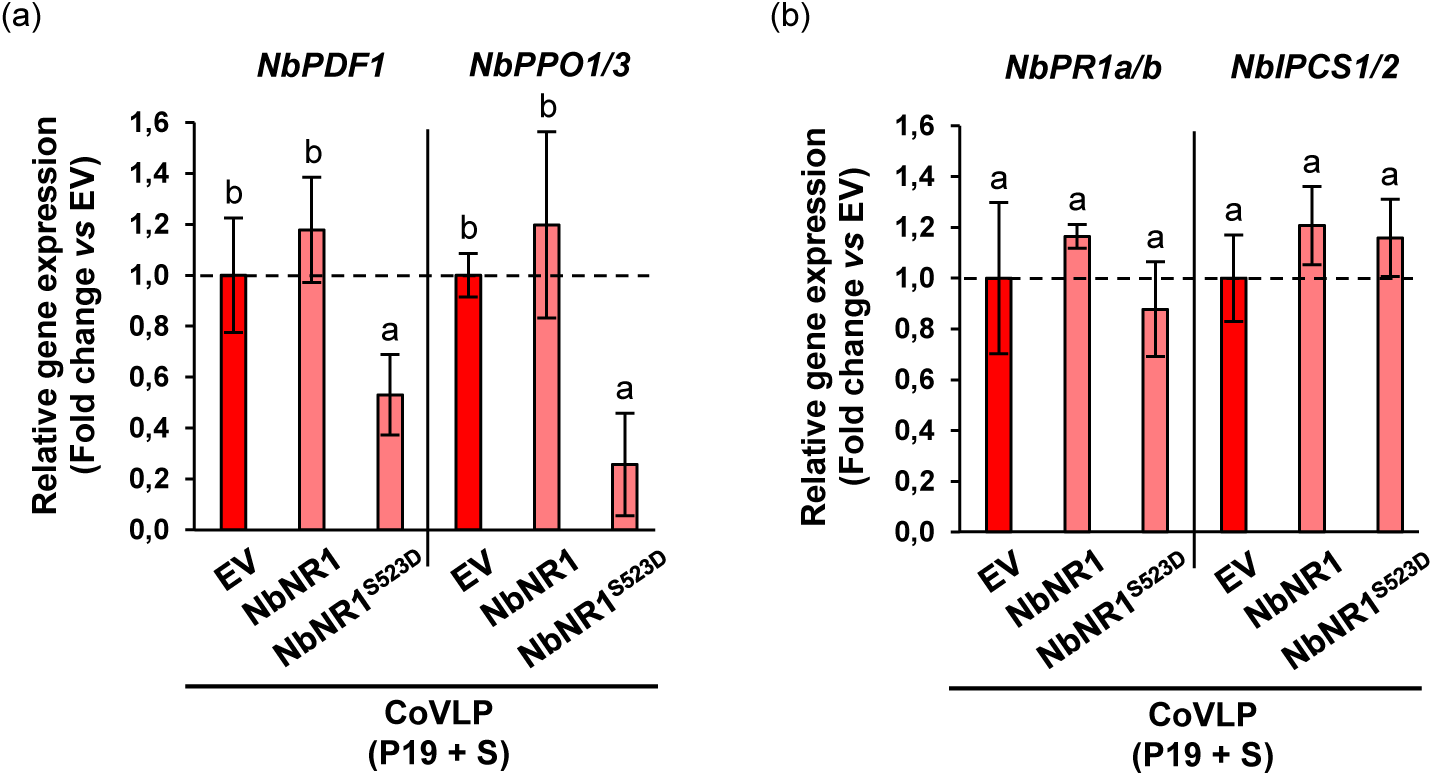
NbNR1^S523D^ does not affect expression of all defense genes. (a) Relative expression of CoVLP-induced genes as measured by RTqPCR. As observed for other defense genes (Figure 8e), expression levels of *NbPDF1* and *NbPPO1/3* were significantly reduced upon co-expression of NbNR1^S523D^. By contrast, expression levels of *NbPR1a*/*b* and *NbIPCS1/2* were unaffected by co-expression of the helper proteins. For each gene, transcript level of CoVLP + EV samples was arbitrarily set at one-fold (dashed line). Error bars reflect the se. Groups that do not share the same letter are statistically different.

**Figure S3.**
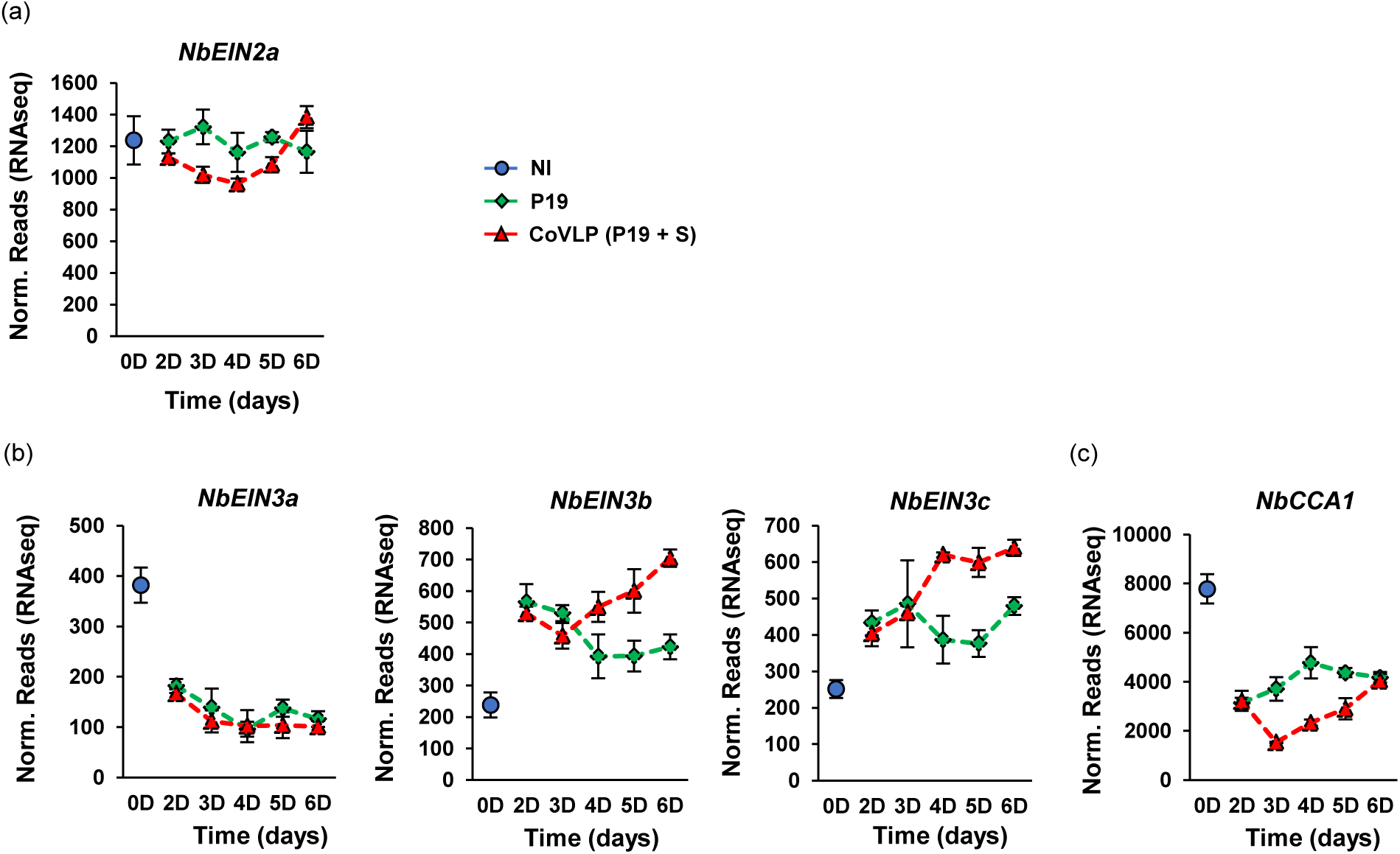
Expression of other regulatory genes involved in senescence. Expression profiles of ethylene signalling genes that encode proteins homologous to Arabidopsis AtEIN2 (a) or AtEIN3 (b). Expression profile of the *MYB*-related gene *NbCCA1* is also shown (c). For each time point in days (D) post-infiltration, RNAseq results are expressed in normalized (norm.) numbers of reads ± sd. Conditions are as follows: NI: non-infiltrated samples (blue); P19: agroinfiltrated samples expressing P19 only (green); CoVLP: agroinfiltrated samples co-expressing P19 and recombinant S protein (red).

**Table S1.** Expression of plant genes as measured by RNAseq on P19 and CoVLP samples harvested at various time points.

**Table S2.** CPs showing enhanced accumulation as measured by iTRAQ proteomics on P19 and CoVLP samples at 6 DPI.

